# Context matters: A meta-analysis of the variable impact of transgenerational and developmental plasticity on responses to stress

**DOI:** 10.1101/2025.08.13.670161

**Authors:** Isabelle P. Neylan, Ana L. Salgado, Rujuta V. Vaidya, Cullen Guillory, Morgan W. Kelly

**Author notes:** Corresponding author: Isabelle Neylan.

## Abstract

1. Understanding organisms’ abilities to adapt and acclimate to stressors in their environments is essential for predicting the distributions and persistence of species and populations during environmental change. Beyond genetic adaptation, prior experiences with a given stressor across life stages can dictate how an individual will fare when exposed to that stressor again. There is now a robust literature on the impacts of parental experiences on offspring traits (transgenerational plasticity), plus an even broader literature on the carry-over effects of early-life experience on phenotypic outcomes (within-generational or developmental plasticity); however, less is understood about the relative strengths of these two forms of plasticity and how they interact to shape stress tolerance.
2. We explored these questions by conducting a meta-analysis of peer-reviewed studies that tested both within- and transgenerational effects of naturally occurring environmental stressors. In particular, we explored contextual moderators or predictor variables including the traits measured, the type of stressor, taxonomic group, and organismal life history traits to elucidate patterns and develop a predictive framework for when we should expect to see effects of transgenerational plasticity, within-generational plasticity, both, or neither.
3. We found that there was not a strong or consistent directional effect of either parental or early-life exposure on subsequent offspring traits. Instead, experimental context (what stressor was used and what traits were measured) as well as biological context (taxonomy, life history traits) were important predictors for understanding the strength and direction of plasticity. We found several contexts where there were meaningful effects of parental and early-life stress exposure and where there was evidence that these may interact to impact phenotypic and fitness outcomes.
4. Our study highlights the need for careful consideration of context when exploring patterns of plasticity. We hope to underscore the need for additional, fully factorial studies that measure the interaction between these forms of plasticity across a variety of systems and stressors to better understand how stress may carry forward across life stages and generations.

## INTRODUCTION

Understanding how organisms respond to environmental stressors is essential to predicting population persistence and is becoming increasingly important in our rapidly changing world. In order to survive, organisms can move or migrate away from stressors, they can adapt via genetic responses to selection over multiple generations, or they may acclimate via phenotypic plasticity, which allows them to more rapidly change their phenotype in response to the environment (Day & Bonduriansky, 2011; Schlichting & Pigliucci, 1998). The impacts of stress may be felt across life stages and may even carry forward to impact offspring. These within and across generation carryover effects may be beneficial if they cue hardening responses that protect against future exposures, or they may be detrimental if the negative impacts carry over to make organisms more vulnerable to future stresses (Bonduriansky et al., 2012; Engqvist & Reinhold, 2016; Uller et al., 2013). We have limited understanding of the long-term outcomes of stress exposure, how plasticity may be impacting patterns within and across generations, and ultimately whether organisms are able to acclimate and adapt, and this is an important gap to address in order to understand responses to environmental stress and global change (Bell & Hellmann, 2019; Donelan et al., 2020; Donelson et al., 2018).

The classic interpretation of plasticity is the ability to alter phenotype in response to the environment, without a genetic change, within a generation (Schlichting & Pigliucci, 1998). Generally, plastic strategies are favoured when the environment is heterogenous, cues about the environment are reliable, and the cost of plasticity is low (Auld et al., 2010; DeWitt et al., 1998; Murren et al., 2015). One particular form of within-generational plasticity is developmental plasticity, which occurs when the environment during an early life stage of an organism (classically seedlings, larvae, or embryos) impacts their phenotype later in life, often irreversibly (Arthur, 2000; Stager et al., 2024; West-Eberhard, 2003). Developmental plasticity has been shown to have strong positive impacts on fitness and there is a robust literature examining its impacts on evolutionary trajectories (Gilbert et al., 2015; Uller et al., 2020, 2024; West-Eberhard, 2003). A more recent body of literature has begun to investigate another form of plasticity where the environment or experiences of parents can impact the phenotype of offspring without genetic change, known classically as maternal effects or more recently as transgenerational plasticity (TGP) (Holeski et al., 2012; Mousseau & Fox, 1998; Salinas et al., 2013). Like within-generational plasticity, there is increasing evidence that TGP can be adaptive or maladaptive and can have impacts on population persistence and evolutionary trajectories (Bonduriansky & Day, 2009; Laland et al., 2015).

There is clear evidence for the importance of both of these forms of plasticity in shaping organismal traits, yet their combined effects are much less well investigated, despite a high likelihood that these two forms of plasticity are interacting in potentially impactful ways (Clement et al., 2023; McNamara et al., 2016; Uller, 2008). A parent’s pre-reproductive experience with stress is likely to be highly temporally (and potentially spatially) correlated with the early-life stages of their offspring. Early life may be a time when organisms benefit most from information provided by parents as they themselves may lack the experience or ability to process information from their own environment (Stamps & Frankenhuis, 2016; Stamps & Krishnan, 2017). On the other hand, stress may weaken parents’ ability to provide essential resources to their offspring leaving them at a disadvantage as they face their own environmental conditions (Uller & Pen, 2011). In the same way, early life stages in many systems are critical, vulnerable windows where organisms are hit hardest by stressors. Effects of stress may carry forward with negative impacts, but early exposure could also harden or prime organisms for future exposures leading to increased tolerance both of which may impact how adult parents go on to prime their own offspring (Mueller, 2018; Przeslawski et al., 2015).

Previous meta-analyses and literature syntheses investigating the strength of TGP have demonstrated that effect sizes are often weak or highly context dependent (Uller, 2008) although not all studies agree (Yin et al., 2019). These mixed results imply that additional context may be necessary to predict patterns of TGP. We hypothesize that one overlooked context in previous TGP studies is the inclusion of the offspring’s early-life environment and its impacts on subsequent trait values and fitness later in life. Carry over effects from both parental experience and early-life experiences are likely to impact offspring phenotypes, but the interactions between these two factors have yet to be explored systematically. In particular, it remains unclear how an organism should respond when receiving contradictory cues from parents and their own developmental experience (e.g., if parents experience stress but the offspring itself does not or vice versa) a potentially underexplored form of parent-offspring conflict (Uller, 2008). A related but distinct question is how a parent’s developmental environment may interact with the environment it experiences as an adult to impact transmission of information to its offspring, or put another way, how a parent’s developmental plasticity may impact its TGP and therefore the traits of its offspring. While the adult environment is more likely to be temporally correlated with the offspring environment, a parent’s early-life experience could serve as a more relevant cue for offspring during the same life phase. This may be a particularly important consideration for species with biphasic development where larvae and adults can experience drastically different selection pressures (Minelli & Fusco, 2010; Pechenik, 2006; Sherratt et al., 2017).

We conducted a meta-analysis using studies that experimentally manipulated both parental and offspring environments by factorially exposing individuals to naturally occurring control and stressful conditions. Our primary goal was to investigate the relative magnitude of stress carry-over effects from developmental plasticity versus TGP on offspring functional traits and performance, and whether there was an interaction between these two forms of plasticity. We found two experimental designs that addressed this question. The first approach used studies that manipulated parental experience and early-life offspring experience and then measured offspring traits related to performance later in life, allowing us to measure effect sizes for TGP, developmental plasticity, and their interaction (Design A). The second approach used studies that manipulated the stress exposure of parents in both early-life and later in life and then measured traits in their offspring, exploring the effect of parental developmental plasticity on transgenerational carryover (Design B). We used a broad meta-analytic approach for both designs to test for generalized patterns and performed a series of meta-regressions using treatment moderators to better understand how TGP and developmental plasticity individually and interactively impacted offspring traits. We expected to observe a mixture of responses to stress, which would contribute to an overall lack of mean effects in either direction when looking across contexts (i.e., in the meta-analysis). This result would align with the results of previous TGP meta-analyses (Byrne et al., 2020; MacLeod et al., 2022; Uller et al., 2013). We also predicted that TGP and developmental plasticity would interactively affect offspring phenotypes and that the treatment combinations where cues matched across time points (e.g., if parents and offspring both experienced control or stress conditions) would elicit the strongest responses.

Based on previous meta-analyses and the empirical literature, we also predicted that the strength of plasticity would differ across contexts. We hypothesized that organismal life history in particular would be an important consideration that would dictate the degree of correlation between parental and early-life conditions. Life history traits that increase the match between conditions within and across life stages (therefore making the environment more predictable) should strengthen selection for adaptive plastic responses to environmental cues (Dury & Wade, 2020; Stamps & Krishnan, 2014). For example, in species with offspring that travel shorter distances away from parents through life history strategies (e.g., internal gestation versus external gamete release) and lack of distinct life phases (e.g., direct versus biphasic development) we should observe increased plasticity and carryover effects. Therefore, our secondary aim was to further explore the context surrounding these patterns to develop a predictive framework for when we should expect to see each form of plasticity. With that goal, we ran additional meta-regressions using contextual moderators that served as predictor variables and captured experimental context (type of stressor being studied, the type of trait being measured in the offspring, and life stage when that trait was measured) as well as biological context (taxonomy, life history traits, and developmental modes). Overall, we predicted that including developmental plasticity in our examination of TGP would illuminate additional patterns suggesting that both forms of plasticity are worth considering in tandem. Finally, we hoped to use our systematic literature review to highlight gaps in the empirical work to inform future research and make suggestions for best practices moving forward.

## MATERIALS & METHODS

### Literature search and study selection criteria

Our goal was to obtain a representative and relatively complete sample of experimental studies that included measurements of both parental and within-generational stress effects and their interactions. Given that TGP studies are a more recent addition to the plasticity literature (Bossdorf et al., 2008; Uller et al., 2013) and are more likely to contain key phrases in their titles, key words, and abstracts (i.e., transgenerational or intergenerational plasticity or effects, parental/maternal/paternal effects, non-genetic inheritance), we decided to begin our literature search by collecting all TGP studies and then screening this larger pool of studies to see which included within-generational plasticity as a later part of our selection criteria. We accessed *Web of Science* and *Scopus* databases on April 17, 2024 and used tailored search strings (see Supplement Methods) and no timespan limit to collect all studies on TGP. This initial search produced 11,181 unique studies (Fig. S1).

For our first round of screening, we kept all TGP papers that experimentally manipulated parent and offspring conditions and measured at least one trait in the offspring. At this stage, we rejected any studies involving humans or human health, agricultural and domesticated species, and stressors that would not occur naturally (e.g., epigenetic manipulation, pollutants, or endocrine disruptors) (see Supplemental Methods for full description of our rejection criteria). Ultimately, this produced 1,803 studies which measured TGP in response to naturally occurring stressors; these were included in further screening (Fig. S1). Our second round of screening aimed to find all papers that included both TGP and developmental plasticity. We retrieved the records for each study and read the Methods sections to determine whether they met our inclusion criteria and fit into one of two designs. Our broad criteria included studies that performed an experimental manipulation of parents and offspring using the same stressor across generations in a factorial design.

### Selection criteria for Design A: Parental and early-life offspring stress and their interaction

Our first design (Design A; Fig. 1a) allowed us to examine the potential carry-over effects of parental stress, early-life offspring stress, and their interaction on subsequent trait measures in the offspring. We define “developmental plasticity” for Design A as a stressor applied to the offspring at one life stage with a trait measured at a later life stage or with some time passing between treatment and measurement if all offspring were measured in the same environment. For example, offspring exposed to either control or heat stress, then allowed a recovery period before all being exposed to heat stress again to calculate thermal tolerance would be included (and would be considered the design shown in Fig. S2b). As a counter example, a study where offspring are exposed to control or heat stress and then placed in a respiration chamber at their respective treatment temperatures would not be included as this would not allow us to untangle the role of developmental experience from later-life offspring experience. We required there to be manipulation of parents and early-life stage of offspring using the same stressor in a fully factorial design allowing for all treatment combinations. However, the final stage where offspring traits are measured was not required to be factorial, although this information was retained and became the offspring treatment moderator (See Fig. S2 for examples of incomplete factorial designs that were included).

**Figure 1.**
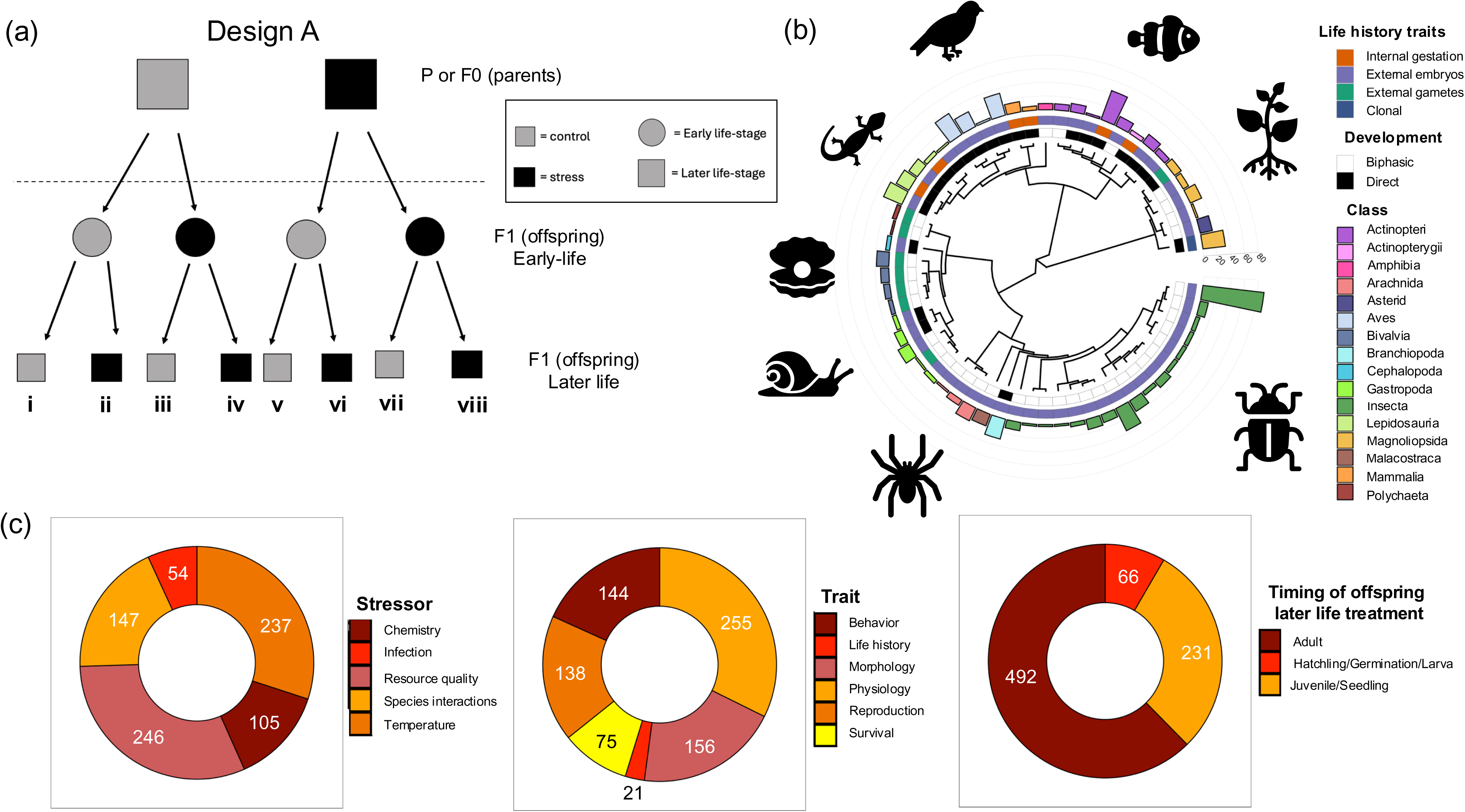
Overview of Design A. (a) A schematic of a fully factorial design of included studies. Gray indicates control conditions, black indicates stress conditions while squares indicate a later life stage and circles indicate an early life stage. Roman numerals are used to differentiate the groups (b) A phylogenetic tree created from the list of species included in our study. The multi-colored bars represent the number of effect sizes for each class of organism (with a color key on the right). The inner bands represent life history traits (colored) and developmental traits (black and white). (c) Graphical representations of the number of effect sizes for our experimental moderators (from left to right stressor type, trait type, and the timing of offspring treatment in later life).

### Selection criteria for Design B: Timing of parental stress and transgenerational carryover

Our second design (Design B; Fig. 2a) allowed us to ask whether the timing of parental stress (early-life or adulthood or their interaction) impacts transgenerational effects on offspring traits. For this design, we required there to be some experimental manipulation of parents with the same stressor used in early-life and adulthood in a fully factorial design allowing for all treatment combinations. Offspring treatments were not required to be factorial, although this information was retained and became the offspring treatment moderator (See Fig. S3 for examples of incomplete factorial designs that were included; See Supplemental Methods for additional details).

**Figure 2.**
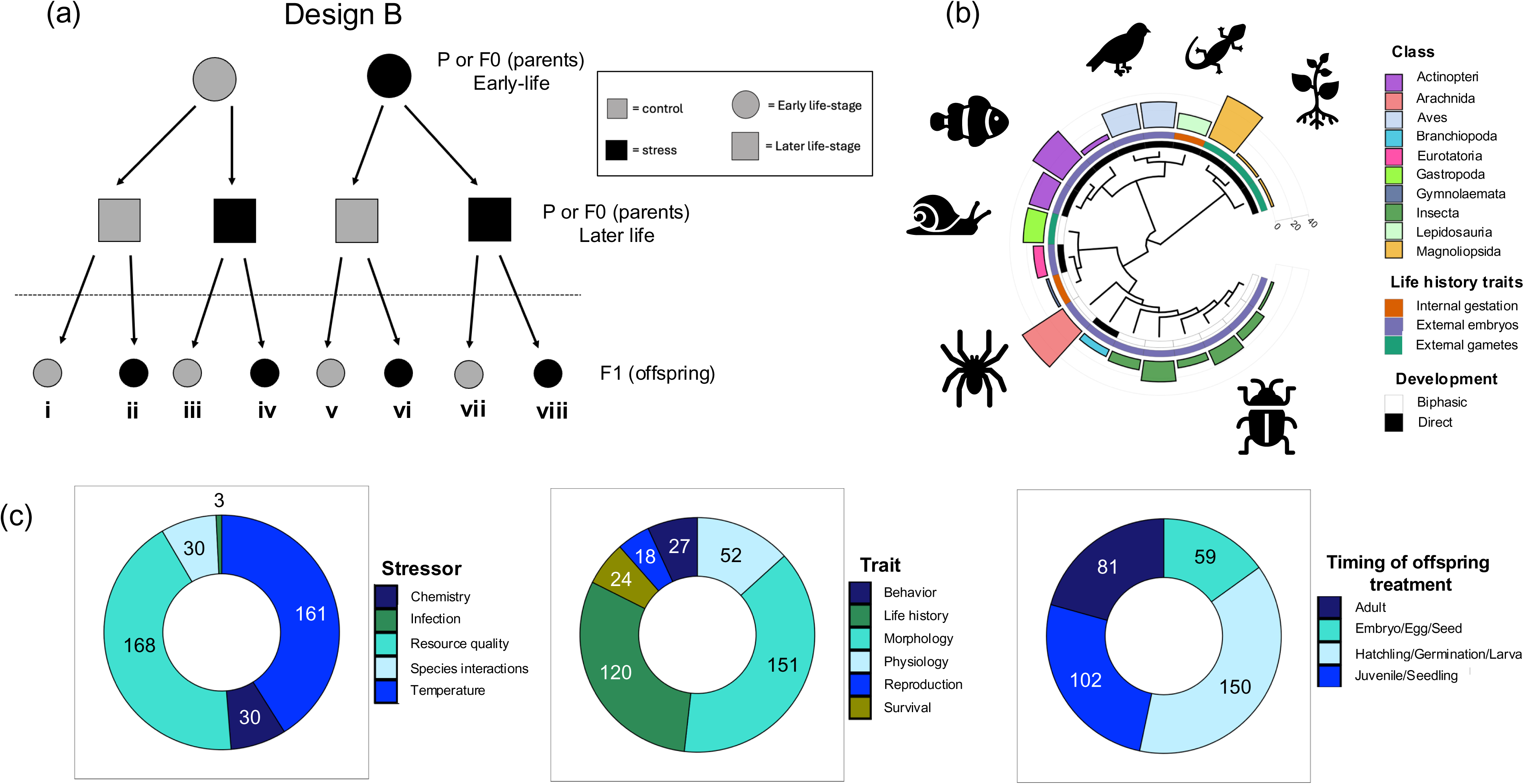
Overview of Design B. (a) A schematic of a fully factorial design of included studies. Gray indicates control conditions, black indicates stress conditions while squares indicate a later life stage and circles indicate an early life stage. Roman numerals differentiate groups. (b) A phylogenetic tree created from the list of species included in our study. The multi-colored bars represent the number of effect sizes for each class of organism (with a color key on the right). The inner bands represent life history traits (colored) and developmental traits (black and white). (c) Graphical representations of the number of effect sizes for our experimental moderators (from left to right stressor type, trait tryp, and the timing of offspring treatment).

At the end of the second screening, we were left with 101 studies across our A and B Designs (77 studies used in Design A, 24 studies in Design B) with four studies using a design that allowed us to extract independent data for both designs A and B (Fig. S4). We followed PRISMA guidelines (Preferred Reporting Items for Systematic reviews & Meta-Analyses; O’Dea et al., 2021; Page et al., 2021) and established our *a priori* hypotheses and relevant moderators prior to beginning analyses (see Supplemental Methods for deviations from these plans). All code, data, and additional information is available on Dryad Digital Repository (Neylan et al., 2025) and all included studies are listed in the Data Sources section of our references.

### Data extraction and effect size calculations

For both designs, we collected the mean, sample size, and variance (error terms) associated with offspring traits across all treatment combinations to calculate effect sizes. When possible, we used raw data provided in repositories or provided by authors to calculate these values. If values were presented in a table, we extracted those values. If neither of these options were available, we used the R package *metaDigitise* (Pick et al., 2019) to extract mean and standard error from figures. When a higher trait value was negatively correlated with fitness outcomes (e.g., development time or latency to leave refuge during safe conditions), we multiplied the mean values by negative one to reverse their sign (Uller et al., 2013; Yin et al., 2019). These changes involved 162 effect sizes (34 studies) from Design A and 62 effect sizes (5 studies) from Design B (Table S1).

Along with these data, we also collected relevant information to use as our moderators (or predictor variables) in meta-regressions including experimental information (stressor category - temperature, chemistry (e.g., pH or salinity), resource quality (e.g., quality of food or habitat), species interactions, or infection; type of trait measured - physiological, morphological, life history, survival, or behavior; transmission type – maternal, paternal, or biparental) as well as information about organismal biology (taxonomy - invertebrate, vertebrate, plant; developmental type - biphasic or direct development; life history traits - internal gestation, external embryos, external gametes, or clonal; timing of offspring trait measure - embryo/egg/seed, hatchling/germination/larva, juvenile/seedling, adult). For both taxonomy and trait type, we chose to keep our categories the same or similar to those used by Uller et al. (2013) and Yin et al. (2019) for ease of comparison (with some modification, see Supplemental Methods - Deviations from predetermined analysis plans).

We used the log response ratio (lnRR) to calculate effect sizes as this calculation tends to be more normally distributed and is more robust to scaling discrepancies, which we thought was a reasonable concern given the diversity of traits being measured in our study (Hedges et al., 1999; Nakagawa, Yang, et al., 2023). However, the log response ratio can also be susceptible to bias when sample sizes are small or the mean is close to zero (Lajeunesse, 2015), which is another relevant concern with our dataset, so we applied a correction to account for this possibility in both our log response ratio and its variance (Yin et al. 2019; Equations 1 and 2). In these equations, the subscript 1 refers to the control treatment while the subscript 2 refers to any stress treatment. All effect sizes were calculated as a pairwise comparison between a control treatment and a stress treatment. For data where offspring traits were measured in control conditions, the control (subscript 1) refers to organisms exposed to control conditions for all time points (parental, and early-life offspring for Design A and early-life and adult parent treatments for Design B) while stress applied at any of those time points became a stress treatment (subscript 2). We coded the timing of the stress as a moderator (see the Meta-regressions section below). For each control treatment, the same value was used in three pairwise comparisons (control-control-control (designated as “i” in Fig. 1a and 2a) vs. control-stress-control (iii), stress-control-control (v), stress-stress-control (vii)). For offspring traits measured in stressful conditions, the control used in the pairwise comparison was based on offspring also measured in stress conditions (i.e., control-control-stress (ii) *not* control-control-control (i)). This control condition was likewise used in three pairwise comparisons (control-control-stress (ii) vs. control-stress-stress (iv), stress-control-stress (vi), stress-stress-stress (viii)). This approach was chosen so that studies that were not fully factorial when offspring traits are measured (offspring measured only in control or only in stress conditions, see Fig. S2 and S3) could be included. The condition during offspring trait measurement is included as a moderator to account for any differences between measurements occurring under control or stress conditions, but it is important to note that we cannot directly compare the effect of stress at the time of offspring trait measurement on those traits due to this design. As this was not the goal of our study and we are more interested in understanding how a stressful context may affect the impacts of stress experienced at previous timepoints, we considered this an appropriate approach to our pairwise comparisons.

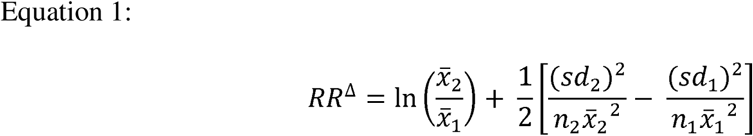

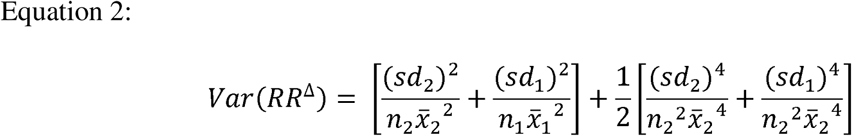

Several of our studies in both Design A and B had a mean or standard deviation of zero, which our effect size equations cannot accommodate. We added a continuous correction of 0.5 to these specific incidences as the literature suggests this is a simple but robust way to handle these cases (Sweeting et al., 2004; Weber et al., 2020) and is preferred to dropping the studies completely (Zabriskie et al., 2024). We also had several studies that were missing error terms in our Design A analysis. To account for these cases, we imputed the standard deviations using a weighted covariance calculated using all data that had standard deviation terms to estimate the missing values (Koricheva et al., 2013; Nakagawa, Noble, et al., 2023). We examined the impact of both the imputations and continuous corrections on our data in our sensitivity analysis.

### Meta-analysis

All analyses were conducted in R (version 4.4.2). Due to various sources of non-independence in our data, we ran multilevel meta-analytic models and meta-regressions using the package *metafor* and the *rma.mv* function (Kim et al., 2020). We ran models on our data from Design A and Design B separately but used the same approach for both designs. For both meta-analytic models, we used t-distributions when calculating confidence intervals and test statistics and fit our models with a restricted maximum likelihood estimator (Nakagawa, Yang, et al., 2023). One source of non-independence in our analysis came from multiple effect sizes being calculated from each study and multiple comparisons made with single effect sizes. To account for this, we included both study identity and an observation-level effect size identity as random effects allowing us to calculate within- and between-study heterogeneity. In addition, we fit a variance-covariance matrix to capture our non-zero covariance due to the fact that multiple traits were measured from the same set of individuals within studies and set rho to 0.5 (which was confirmed to be a robust value in subsequent sensitivity analyses, see below; (Nakagawa, Yang, et al., 2023; Viechtbauer, 2010). One final source of non-independence came from the fact that the studies used a wide range of organisms with varying levels of phylogenetic relatedness. We constructed a phylogenetic matrix to account for the level of relatedness across each species using the *rotl* and *ape* packages (Michonneau et al., 2016; Paradis & Schliep, 2019) and constructed a phylogenetic tree to create a correlation matrix. Branch lengths were computed using Grafen’s method and polytomies were resolved at random. One species had to be manually added to the tree in the Design A analysis (*Siphonaria australis*). We also used the *taxize* package (Chamberlain & Szöcs, 2013) to assign each species with their taxonomic information including phylum, class, order, and family.

When conducting model selection with our random effects, we found that for both Designs A and B the phylogenetic matrix explained very little of the variance and did not improve model fit. Just including genus and species without accounting for relatedness produced the best fitted model (as determined by AIC score comparisons; Table S2). We therefore moved forward with study identity, effect size identity, and species identity as our random effects along with our variance-covariance matrix for both designs. Using this model construction, we calculated the overall meta-analytic mean and the relative heterogeneity (variance due to differences between studies, not due to sampling variance) and calculated how much each random effect contributed to this heterogeneity for each design.

### Meta-regressions

Addressing our hypotheses required the addition of treatment information and contextual moderators which we explored using a series of meta-regressions. We placed our moderators into two categories. The first were treatment moderators that were integral to interpreting our results in terms of plasticity and stress carry-over. These included the parental treatment, early-life offspring treatment, and final offspring treatment in Design A (Fig. 1a) and early-life parental treatment, later life parental treatment, and offspring treatment in Design B (Fig. 2a) (all coded as either control or stress). We began by running single moderator models for each of the treatment moderators to see if we could detect a signal of transgenerational or developmental plasticity across studies and contexts. For each regression, we visually assessed homogeneity of residual variance and when heteroscedasticity was detected, we used the “HCS” structure with zero covariance from the *rma.mv* function from the *metafor* package (Kim et al., 2020) to account for the variance within moderator categories. This heteroscedasticity was most likely due to our high level of variance in effect size identity (see Results). We then built a more complex model with all three treatment moderators and their interactions, including a three-way interaction, and assessed model fit and subsequent results.

Our second set of moderators provided additional experimental and biological context. These included taxonomy, stressor category, trait type, transmission type, life history traits, developmental traits, and timing of the offspring measurement. We used the same approach and began by modelling each moderator individually and assessing the model for heteroscedasticity as described above. We then ran our multiple moderator regression models. All models included our treatment moderators, as they were necessary to address our hypotheses (e.g., we were interested in how trait type might impact TGP, not just the response to stress more broadly). To help determine the best fit of these complex models and determine the model with the highest explanatory power, we used the *dredge* function in the *MuMIn* package (Barton, 2009) to fit models with combinations of moderators and then used AICc (Akaike information criteria corrected for small sample sizes) to determine the best-fitting models and the relative importance of each moderator.

Unfortunately, our full model with all interactions was too large and complex to dredge in one run, so we dredged multiple models with as many combinations of moderators and interactions as possible and used this information to identify moderators with explanatory power worth keeping in our model. We could not accurately determine model structure or weights from these dredging results so proceeded with additional model selection methods.

We first fit and ran a full model with all treatment and context moderators and all interactions. We then ran a reduced model with all three treatment moderators, but only those context moderators identified as important by our dredging results, plus interactions between all moderators. Finally, we ran a further reduced model, again with all three treatment moderators and context moderators determined by dredging but that only included interactions between all three treatment moderators and each context moderator (i.e., no interactions between context moderators) as this best addressed our hypotheses and maintained adequate sample sizes within interaction categories. This third reduced model proved the best fit for both Design A and B (see Tables S7 for Design A and S12 for Design B for more details).

### Publication bias and sensitivity analyses

We assessed publication bias separately for both Design A and B through visual inspection of funnel plots and by testing the robustness of the meta-analytic models by comparing our results with models that included a publication-bias-corrected effect (the adjusted sampling error) (Nakagawa, Yang, et al., 2023). We also tested for small study effects by adding the uncertainty of effect size into our models (Egger’s regression) (Egger et al., 1997; Nakagawa, Yang, et al., 2023). Finally, we tested for a time lag bias or decline effect by including the centered publication year.

We used several approaches to assess the sensitivity of our models to different sources of variance. We ran a leave-one-out analysis for both Design A and B to determine if any study was considered an outlier or overly influential in our results. In Design B we found one study that could be considered an outlier and therefore ran a meta-analytic model leaving this study out and compared these results to our meta-analytic results including all studies. We followed a similar procedure for determining the potential influence of studies with imputed standard deviations and those where we applied continuous corrections. Finally, we compared model outputs using various values of rho to test the robustness of our variance-covariance matrix construction.

## RESULTS

### Literature overview & gaps

We collected a total of 1,250 effect sizes from 101 studies spanning the years 1993 to 2024 that used 70 unique species. These effect sizes were spread across our two study designs (Design A Fig. 1 and Design B Fig. 2) with Design A having more studies and effect sizes overall (846 effect sizes from 77 studies) compared to Design B (404 from 24 studies). Both designs included a broad distribution across taxonomy, life history and developmental traits, stressors, trait measures, and timing of offspring measures (Fig. 1b,c and Fig. 2b,c). Overall, there were far fewer plant studies than we expected *a priori* given the prevalence of plants in the TGP literature. There were also far fewer papers dealing with infection than we expected given their prevalence during our first screening for TGP papers and the large body of transgenerational immune priming (TGIP) literature.

### Results for Design A: Parental and early-life offspring stress and their interaction

Overall, we did not find a directional effect of stress (experienced across any time point, in parents or in early life, in any combination) on offspring trait values as revealed by our broad meta-analytic model (Fig. 3a; Table S3). This result was due to many studies having effect sizes close to zero, but also multiple strongly positive and negative effect sizes. Heterogeneity was extremely high (*I*^2^ = 99.8%) with 81.4% explained by within-study heterogeneity (effect size variance), less than 1% explained by variation between studies, and 18.4% explained by differences across species (without accounting for phylogenetic relatedness which explained even less variation and was dropped from the model).

**Figure 3.**
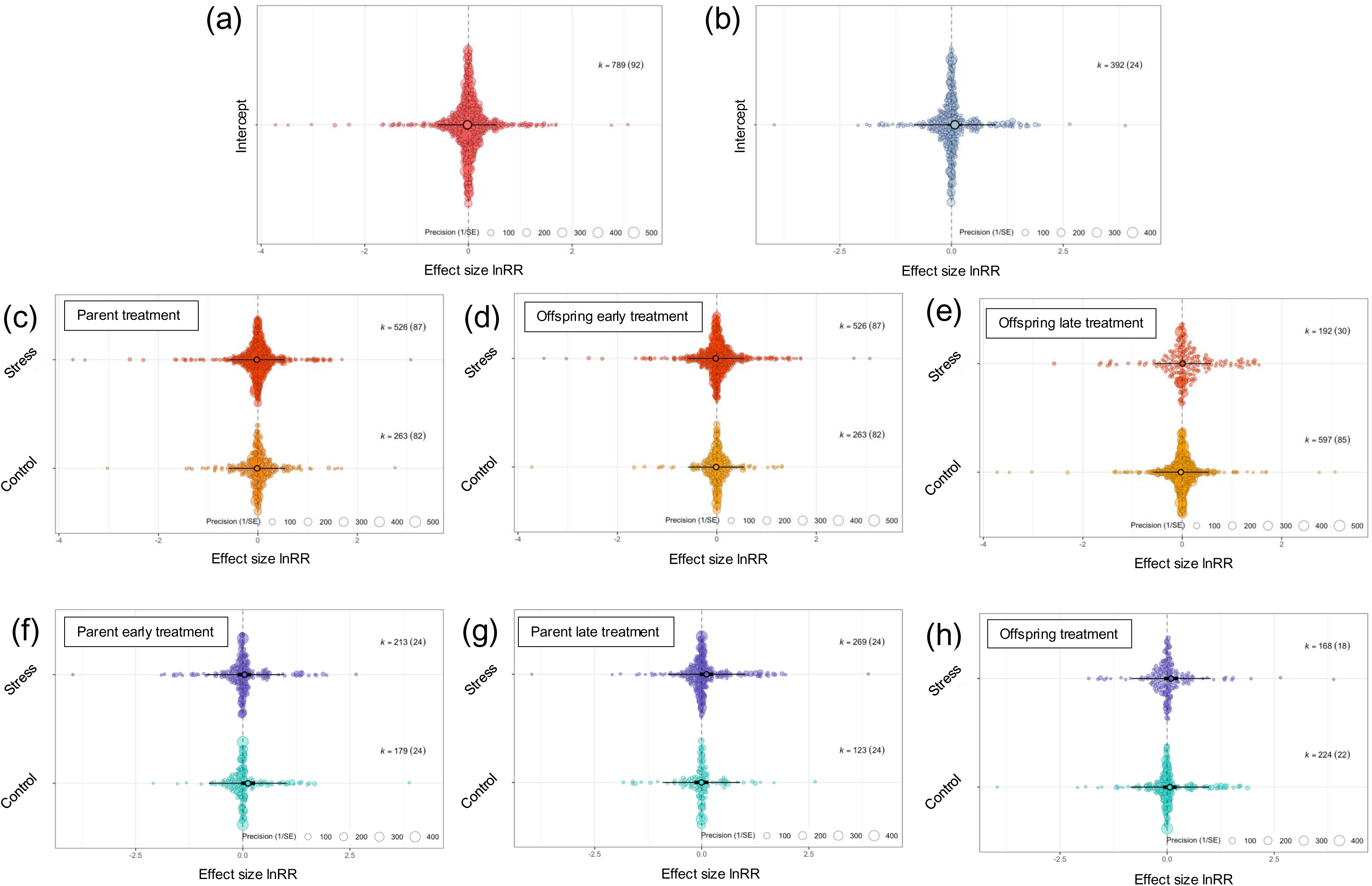
Overall effect of stress on offspring traits in Design A (a) as well as the impact of control versus stressful conditions in parents (c), the early-life stage of offspring (d), and the later life stage of offspring (e). Similarly, the overall effect of stress on offspring traits in Design B (b), as well as the impact of control versus stressful conditions in the early life of parents (f), a later life stage in parents (g), and in offspring (h). Individual colored circles represent effect sizes and are scaled by precision (inverse of standard error). The central thicker circles indicate the mean estimate with thick error bars representing the 95% confidence intervals and thinner bars representing the prediction intervals (note for graphs (a)-(e) the thicker bars are hidden behind the mean estimate circle). K = number of effect sizes (number of studies).

Our meta-regressions using single treatment moderators (Table 1) also revealed no strong patterns and each treatment moderator explained very little of the residual variance (Fig. 3c,d,e; Tables S4) suggesting that neither parental nor developmental effects had a consistent positive or negative effect on later offspring traits. The meta-regression including all three treatment moderators and their interactions similarly provided no significant single moderators or interactions (p-value > 0.2 for all; Fig. S5; Table S5). However, we began to see estimates significantly differ from zero when we included our context moderators. In particular, we found that stressor category and trait type were important explanatory variables (Table 1). Studies that manipulated resource quality had a significantly negative outcome on offspring traits generally (t-value = −2.751, p-value = 0.006) while any stress treatment significantly reduced survival (t-value = −2.345, p-value = 0.019) but increased the means of traits related to life history compared to controls (t-value = 2.823, p-value = 0.005; Fig. 4; Table S4) regardless of whether the stress was experienced by parents or by offspring during early life. These two moderators along with later life offspring treatment were highly heteroscedastic (Table S6) so we used models that accounted for this increased variance and report these corrected results (Table S4).

**Figure 4.**
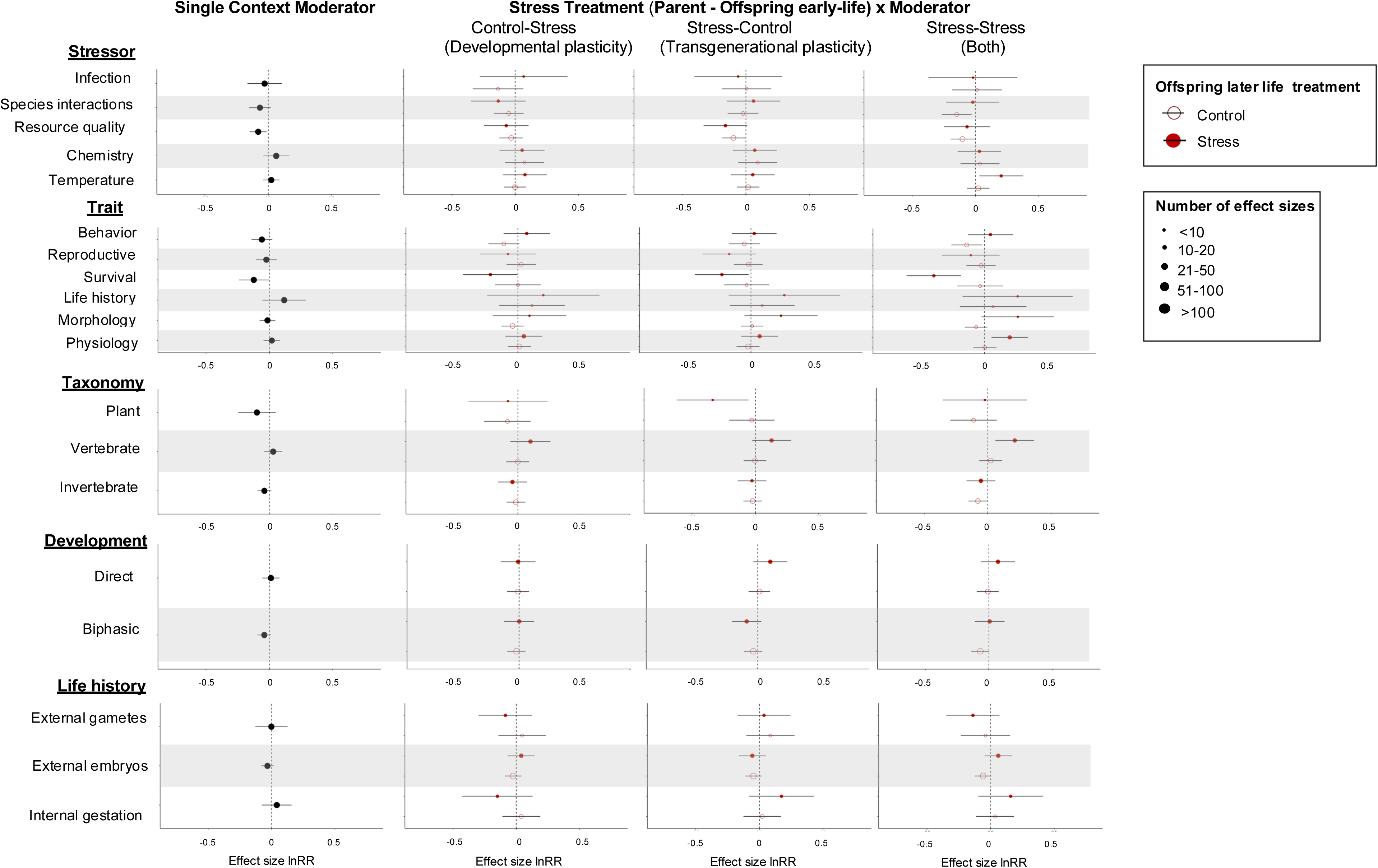
The variation in carryover of stress depending on context for Design A. The first column shows the overall effect of each moderator category on the effect of stress as measured by offspring traits. The context moderators shown here had statistically significant results as a single moderator and/or in interactions with treatment moderators (see Table S4 for full statistical details). The three columns to the right show the strength of transgenerational and developmental plasticity and their interaction in the various contexts (or treatment by context moderator interactions). The graphs in the right three columns are organized by parent-offspring early life treatment combinations (all calculated in comparison to control-control) with the open circles indicating control conditions during the later offspring life stages while the filled circles indicate stressful conditions. The size of the circles represents the number of effect sizes in each of the sub-categories and the error bars represent the 95% confidence intervals of the effect sizes. Note that we removed clonal as a category in development mode as the sample size was too low.

**Table 1.**
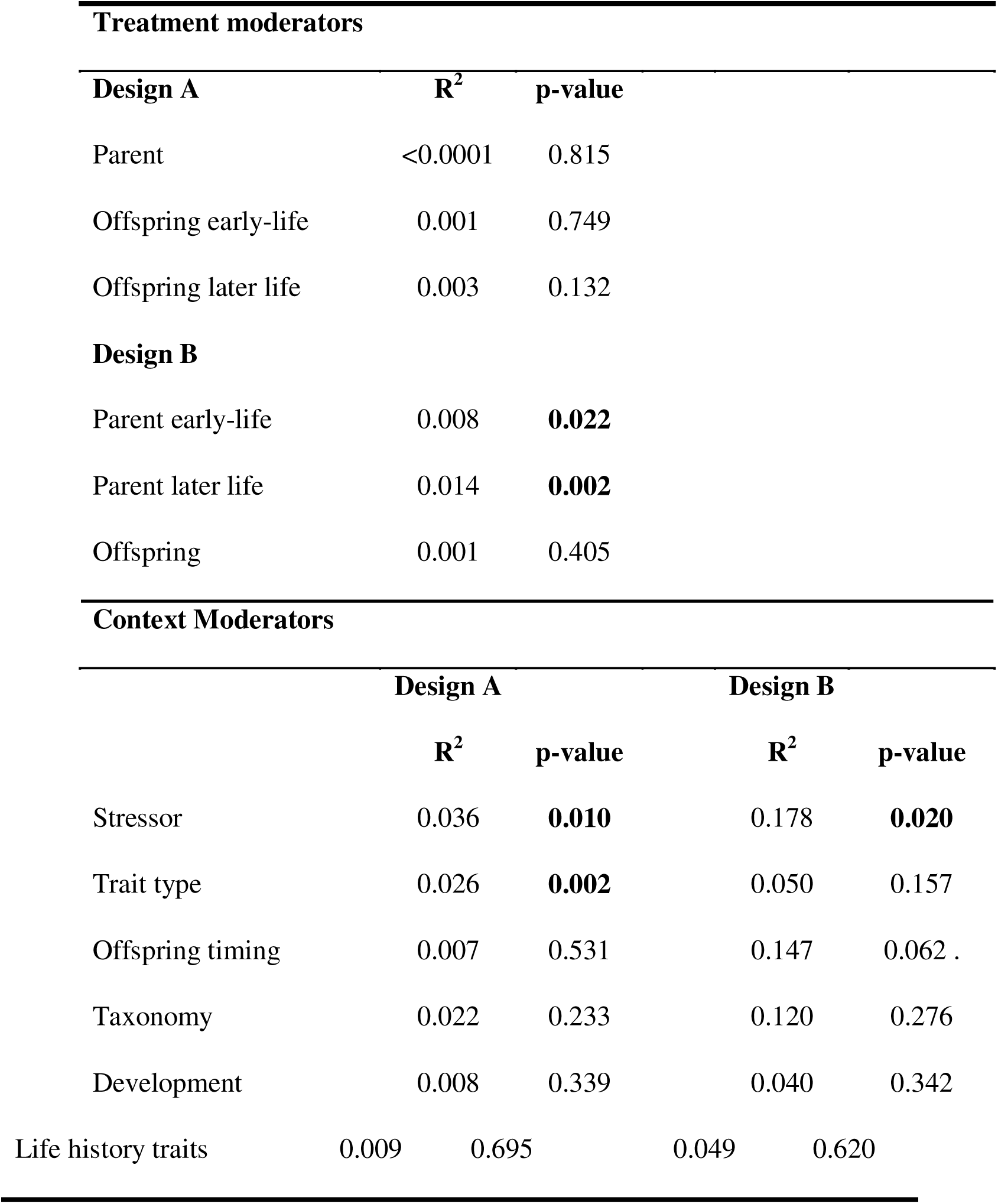
Results from single moderator meta-regressions fitted with treatment and context moderators for both Design A and B. The R^2^ values shown represent marginal R^2^ values. Bolded p-values indicate significance at p < 0.05.

When we explored all moderators and their interactions, we found that the model with the best fit that included our treatment moderators also included stressor category, trait type, taxonomy, development, and life history as context moderators (but dropped transmission type and timing of offspring trait measures) (Table S7). This model found a significant interaction between offspring early-life and offspring later life treatments as well as three-way interactions between these two treatment moderators and trait type, taxonomy, and developmental mode. We also found a three-way interaction between parent treatment, offspring later life treatment, and life history traits.

### Results for Design B: Timing of parental stress and transgenerational carryover

Similar to our results for Design A, we did not find an overall directional effect of stress on offspring traits in our meta-analytic model (Fig. 3b; Table S8). Heterogeneity was again extremely high (*I*^2^= 99.9%) with within-study heterogeneity explaining 46.5% of the variance, differences across species explaining 53.4%, and between study variance again less than 1%. As in Design A, we found no evidence for an effect of our treatment moderators on overall trait means in response to stress (Fig. 3 f, g, h; Table S9). However, our single moderator meta-regressions for parental treatment in both early life and later life proved significant (Table 1; Fig. S6; Table S9) and our meta-regression that included all three treatment moderators and their interactions did significantly explain heterogeneity leftover from our random effects suggesting that these moderators had explanatory power for our B Design (p-value = 0.0297; Table S10).

For context moderators, we found in our single moderator meta-regressions that stress category was a significant explanatory variable with resource quality being the driving category (t-value = 2.020, p-value = 0.044). The trait means in offspring tended to higher when parents were stressed at either early-life or adulthood, but this pattern did not hold when parents experienced sustained resource limitation across both early-life and adulthood (Fig. 5). The model with timing of offspring trait measures was marginally non-significant (Table 1; Table S9; Fig. 5), but we did find that individuals tested as adults had higher mean trait values after experiencing stress than their control counterparts (t-value = 2.230, p-value = 0.002) while we did not detect this pattern when traits were measured at any other time point in offspring (p-value > 0.3 for the rest). This result suggests that the consequences of TGP seem to manifest in adult rather than immature offspring when they encounter the same stressor their parents did. As with Design A, we found multiple moderators lacking homoscedasticity in their residuals, so we ran models that accounted for the heterogeneity (Table S11) and report these corrected results for taxonomy, transmission type, development, life history traits, and trait type (Table S9).

**Figure 5.**
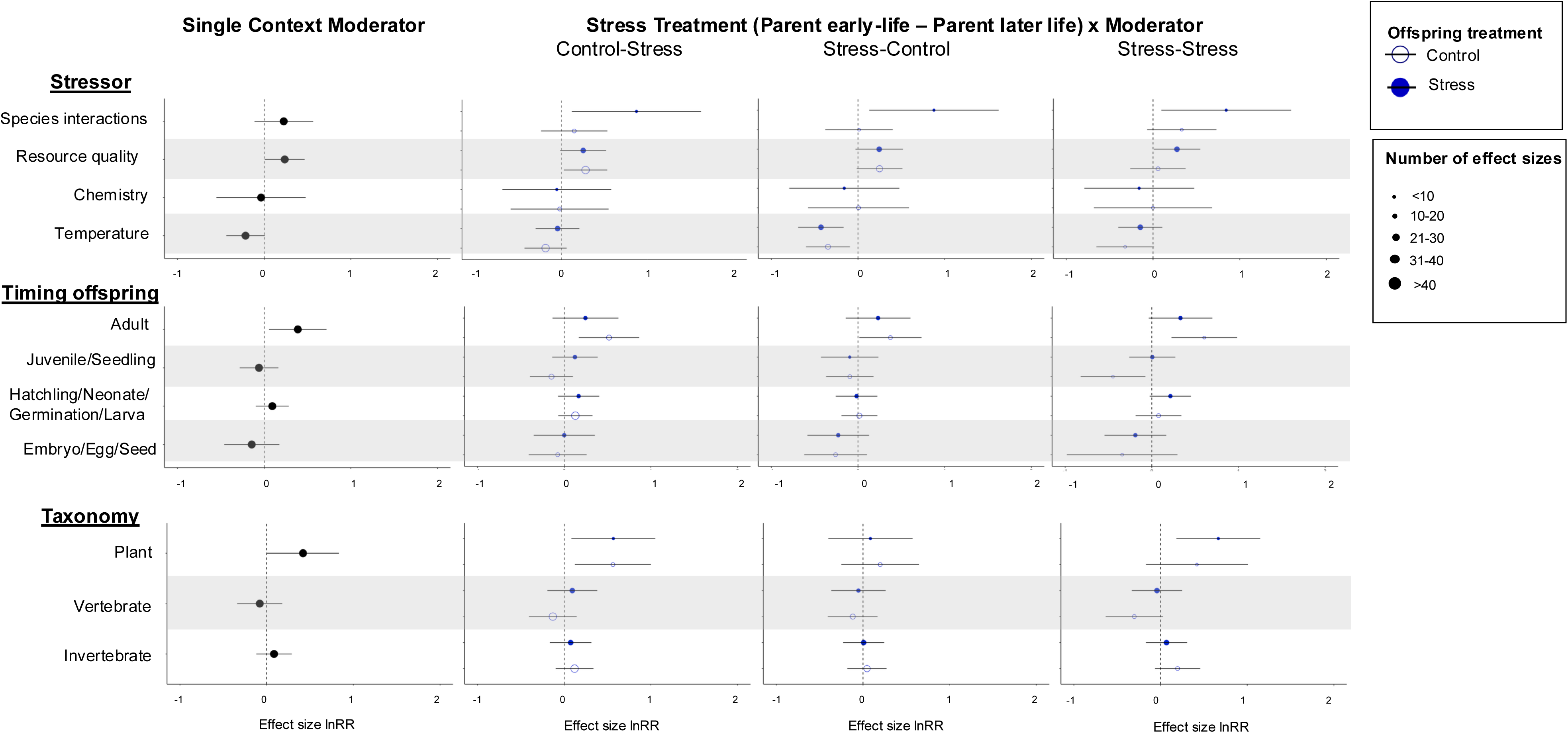
The variation in carryover of stress depending on context for Design B. The first column shows the overall effect of each moderator category on the effect of stress as measured by offspring traits. The context moderators shown here had statistically significant results as a single moderator and/or in interactions with treatment moderators (see Table S9 for full statistical details). The three columns to the right show the strength of transgenerational and developmental plasticity and their interaction in the various contexts (or treatment by context moderator interactions). The graphs in the right three columns are organized by parental treatment combinations (all calculated in comparison to control-control) with the open circles indicating control conditions for offspring while the filled circles indicate stressful conditions. The size of the circles represents the number of effect sizes in each of the sub-categories and the error bars represent the 95% confidence intervals of the effect sizes. Note that we removed infection as a stressor category as the sample size was too low.

Finally, we investigated all treatment and context moderators and their interactions and found that the best fitting model that included all treatment moderators also included timing of offspring trait measures, stressor category, and taxonomy (and excluded life history traits, transmission type, and trait type) (Table S12). Using this model, we found a significant interaction between early-life parent treatment and later life parent treatment as well as three-way interactions with offspring treatment, stressor category, and timing of offspring trait measures while taxonomy interacted with later parental treatment.

### Bias and sensitivity results

Overall, we did not see evidence for publication bias through visual inspection of our funnel plots (Fig. S7), and we did not see a significant difference between our meta-analytical models and those using a publication-bias-correction in either Design A or B (Table S13). Similarly, we did not find significant evidence for small study effects (Table S14; Fig. S8) or for time-lag bias (Table S15; Fig. S9) in either design. Our results proved robust against the removal of any one study for Design A (Fig. S10). Design B, however, had one study that had the potential to be an outlier (Fig. S11), so we specifically re-ran our meta-analytic model without this study, but found the results to be qualitatively the same and very similar quantitatively so do not believe this study to be overly influential on our dataset (Table S16). We also re-ran our analytical models without effect sizes that required imputing standard deviations for Design A (Table S17) and those that required a continuous correction (Design A Table S18; Design B Table S19). We found that none of these models provided dramatically different results and qualitatively the outputs remained the same. Finally, we found that both designs had variance-covariance matrices that were robust to changes in the correlation value of rho (Table S20 and S21).

## DISCUSSION

We conducted a meta-analysis to examine how developmental plasticity and TGP influence offspring traits, focusing on their relative strengths, interactions, and context-dependent effects. We expected to observe a mix of responses to stress, with stronger effects when parental and offspring stress cues matched. Additionally, we predicted that timing, life history traits, and experimental conditions would significantly influence plasticity outcomes, shaping a predictive framework for future research.

### Main findings - does stress carry forward?

Overall, we found little evidence of consistent parental or developmental effects on subsequent offspring functional traits or performance when looking at the data as a whole, but we were able to detect complex plasticity and carry-over patterns when contextual moderators were considered. Our two meta-analytic models (Design A and B) did not show evidence of stress at any life stage impacting offspring traits in a positive or negative direction (Fig. 3a, b). Similarly, we did not find significant effects in any of the treatment moderators in our single moderator meta-regressions or meta-regressions including their interactions (Fig. 3c-h). These results suggest that the impacts of stress are not generalizable or detectable when the data are taken in aggregate.

Importantly, our data were highly heterogeneous, with large confidence intervals and high *I*^2^ values and much of the variation residing within studies rather than between studies. The lack of clear patterns may be due to the amount of diversity covered in this meta-analysis, including taxonomic and biological diversity as well as a variety of stressors and experimental approaches with multiple traits often measured within the same experiment. It is worth noting that in these broad analyses we consider performance traits (survival, reproduction, life history) along with functional traits (morphology, behavior, physiology) which may have contributed to this large level of heterogeneity and underscore the importance of including trait moderators in our meta-regressions (Davidson et al., 2011). The effect sizes were quite variable in both magnitude and direction with many small effect sizes centered around zero, but many also ranging from strongly positive to strongly negative. These results confirm our initial hypothesis that we would see a mixture of plasticity responses across contexts with some studies finding significant adaptive plasticity or maladaptive carry-over responses while others found effects that were more subtle or non-significant (Uller et al. 2013; Bell & Hellmann 2019; Byrne et al. 2020).

Adding in our context moderatos provided a more nuanced picture. In our single moderator analyses, we found that across Design A and B the type of stressor was an important explanatory variable (Table 1; Fig. 4 and 5). Our more complex models show evidence that there is in fact parental and developmental plasticity occurring, but only in certain contexts. Even within a given category, the direction of effects can differ. For example, in Design B, resource limitation stress experienced by parents positively impacted offspring traits (evidence for parental priming) while temperature stress in the parental generation decreased offspring performance (evidence of negative carry-over) especially when parents were only exposed in early-life and not again in adulthood (Fig. 5; Table S9). These results are aligned with our second hypothesis, that TGP and developmental plasticity would interact, but the weakness of these overall effects and the specificity were not what we had predicted *a priori*.

Our results align with previous meta-analyses of TGP that found weak or mixed effects of parental experience on offspring traits (Byrne et al., 2020; MacLeod et al., 2022; Uller et al., 2013). For two of these studies, the focus was on a specific taxonomic group (Byrne et al. 2019 marine invertebrates; aligning with our lack of strong patterns in invertebrates) or stressor (MacLeod et al. 2022 predator-prey interactions; agreeing with our lack of results for species interactions). Yin et al. (2019) and Uller et al. (2013) both used a wide diversity of studies like our analysis, but they differ in their conclusions. While Uller et al. found weak evidence for anticipatory parental effects Yin et al. found an overall positive or adaptive effect of TGP. Our broader results align more closely with Uller et al. in that we do not detect a strong, broad TGP effect. However, Uller et al. did not detect patterns even with contextual moderators while we were able to reveal patterns of TGP. This difference may be due to the number of additional studies that have been published since 2013, which also include a broader diversity of species and contexts. Yin et al. did detect contextual differences in similar categories as our study, but we did not find strong generalizable results (although we did find evidence for TGP effects in Design B). These differences could be due to a number of factors, but in particular our studies differ in that we have focused exclusively on studies using natural and clearly definable stressors while Yin et al. and Uller et al. do not, we have limited the number of TGP studies to those that also include developmental plasticity in a fully factorial design (significantly reducing the number of included studies), and there were some differences in our model construction and analyses. Meta-analyses specifically focusing on developmental plasticity tended to have larger effects than we found in our study (Albecker et al., 2023; Noble et al., 2018; Pottier et al., 2022) although it is worth noting that all three of these studies only explore either one taxonomic group (Albecker et al. Anurans; Nobel et al. Reptiles) or one type of stressor (Pottier et al. and Nobel et al. temperature). This study is the first to our knowledge that combines both TGP and developmental plasticity across a diversity of study systems and experimental contexts. Our findings demonstrate that plasticity is highly context dependent. Importantly, also we observed no relationship between the existence of TGP and the existence of developmental plasticity across systems. Studies which observed significant effects of TGP were no more likely to also observe developmental plasticity than those that did not observe TGP. The reverse was also true with studies that observed developmental plasticity being no more likely to observe TGP than those that did not.

### Design A - How does parental and early-life offspring stress carry forward and is there an interaction between TGP and developmental plasticity?

In the aggregate, we did not find strong evidence for either parental effects (TGP) or developmental plasticity except under certain biological and experimental contexts (Fig. 3a, c-e; Fig. 4). We also did not find any evidence for an interactive effect between parental and early-life offspring treatments (a TGP and developmental plasticity interaction). As we predicted, including additional moderators impacted our results; However, this evidence is subtle and highly context dependent.

Experimental design factors (both stressor category and trait) proved to be important predictors. Stress, in particular reduced resource quality, negatively impacted offspring performance regardless of when the exposure occurred. Interestingly, survival and life history traits were the most impacted by stress, although in opposite directions with survival reduced in response to stress and life history traits positively impacted (Table S4; Fig. 4). These results make some intuitive sense as survival directly relates to fitness outcomes and faster development time (coded in our study as adaptive) has been linked to stress responses in multiple systems (van der Linden et al. 2018; Crespi et al. 2012). Shorter development times and therefore earlier reproduction is a common form of bet hedging in stressful environments to ensure that reproduction happens if survival is not guaranteed. These faster development times often incur trade-offs such as reduced fecundity or lower offspring quality, so while our results for life history traits appear to be a positive outcome of stress, this may in fact be a signal of reduced fitness if parent-offspring conflicts are considered (Crean & Marshall, 2009; Uller, 2008). Selection on viability (survival) has also been shown to more strongly predict anticipatory effects than selection on traits impacting fecundity (Kuijper & Johnstone, 2021).

We also found that both offspring treatments (early life and later life) interacted in multiple biological contexts including trait type, taxonomy, and developmental mode suggesting developmental plasticity did impact organismal traits in these studies, depending on the study system. Our only evidence for parental effects was parental treatment interacting with later life offspring treatment when subdivided by life history traits (external embryos appear to experience possible negative parental effects but only in stressful conditions). These findings indicate to us that developmental plasticity has a stronger effect than TGP and that these two forms of plasticity do not appear to be interacting. While this is not what we predicted, these results do align with growing empirical evidence suggesting a decoupling of within-generational plasticity and TGP where being plastic across one set of life stages does not necessarily mean being plastic across others (Tariel et al., 2020; Walsh et al., 2014).

Theoretical and modelling work have also demonstrated a possible trade-off or compensatory effect between these two plasticity strategies (Clement et al., 2023; Kuijper & Hoyle, 2015; McNamara et al., 2016) with anticipatory parental effects evolving more slowly and having less fitness benefits than within-generational plasticity potentially explaining weaker empirical evidence for TGP (Dury & Wade, 2020; Kronholm, 2022). Additionally, reduced performance of offspring from exposure to stress in parents or in early life may simply indicate a consequence of repeated stress or constraint on performance rather than an active plastic response.

### Design B - How does the timing of parental stress influence transgenerational carryover?

The evidence for parental effects was stronger in Design B although we still failed to detect a consistent effect of stress (Fig. 3b). However, in our single moderator meta-regressions we found that parental stress in both early life and later in life had significant explanatory power (Table 1; Table S9) likely due to the differences in the direction of response between stress and control conditions given that the mean effect size for both treatments did not significantly differ from zero (Fig. 3f, g). These results suggest that there are significant transgenerational effects occurring across contexts and that exposure to stress in both early-life and adulthood can be transmitted to offspring, impacting trait values.

Once again, context proved an important consideration. Like in Design A, studies reducing resource quality across generations and life stages impacted offspring traits, but in Design B we found the opposite pattern with offspring performance generally increasing after parental resource stress, evidence of parental priming (Fig. 5; Table S9). Starvation and low caloric intake have been shown to have strong, multi-generational effects in multiple systems (Paul et al., 2022; Rechavi et al., 2014; Veenendaal et al., 2013). We also found that offspring measured as adults had the strongest evidence for transgenerational effects and these effects were generally positive (Fig. 5; Table S9). This is an unexpected result as we hypothesized that earlier life-stages might show stronger evidence for parental influence given the closer timing between parent and offspring environments. This effect is also strongest in adult offspring that are measured in control conditions while their parents experienced stress, which again does not align with our initial hypothesis that the strongest effects would come from treatments where parent and offspring environments matched. Finally, we found that plants showed the strongest transgenerational pattern with later parental experience positively impacting offspring performance (Fig. 5). This result aligns well with current empirical literature that demonstrates plants are capable of adaptive TGP and are known to faithfully pass down epigenetic marks such as DNA methylation across generations (Galloway & Etterson, 2007; Herman & Sultan, 2016; Sobral et al., 2021).

### Gaps and limitations

Our analysis was limited by the small number of studies and the large amount of heterogeneity that led to a lack of precision in some of our more complex statistical comparisons. By including a wide diversity of taxa, stressors, and traits measured we may be masking some important patterns within a given context and encourage empiricists to continue to explore these questions in their systems of interest. We believe our results are broadly generalizable to systems experiencing natural stressors, but acknowledge that we did not include studies using chemical stressors such as pollutants or using agricultural species which are an appreciable subset of the TGP and developmental plasticity literature. We chose to use naturally occurring stressors and exclude domesticated or artificially selected species as our aim was to better understand plasticity patterns that were part of an evolved response and could inform our understanding of stress responses in nature.

Our statistical approach also meant that we required studies to be fully factorial, which forced us to drop multiple studies that tested relevant questions, but did not follow our design structure (for example 23 studies tested both developmental plasticity and TGP but separately on different individuals or in different experiments and thus could not be included). We also acknowledge that there are limitations to the match/mismatch approach that was used in all of the empirical studies included in this meta-analysis and we may therefore under or over-estimate the adaptive nature of parental effects (Bonduriansky & Crean, 2018; Engqvist & Reinhold, 2016). We urge caution when interpreting our results in terms of adaptive versus maladaptive outcomes, especially given the uncertainty surrounding our complex interactions. However, we believe the structure of our analysis addresses some potential concerns. By tracking stress legacy across multiple time points in a fully factorial manner and incorporating this information in our moderators, we gain insight into when an effect goes beyond just match/mismatch between two treatments during two time points (as in a standard reaction norm) which should help capture the possibility of non-additive or “silver spoon” effects (Engqvist & Reinhold, 2016).

### Future directions

One of the clearest results from our systematic review of the literature was that there is a lack of studies examining the combined effects of developmental plasticity and TGP. Of the 1,803 TGP papers that made it to our second round of screening, we were only able to use 101 of them for this analysis. We believe this is an exciting and fruitful gap to be filled in the empirical literature. In particular, we were only able to find 24 studies that fit our B design, and these results were some of the most intriguing. Considering the timing of parental stress and how multiple exposures across ontogeny may impact the transmission of transgenerational information is an area ripe for additional work. We strongly encourage future studies to incorporate fully factorial designs (Fig. 1a and 2a) in order to explore possible interactions and produce the most robust datasets. Additionally, only four papers were fully factorial and incorporated elements that satisfied both Design A and B. If time and space constraints allow, we believe this is the most robust design to detect both forms of plasticity across generations, life stages and their potential interactions (Fig. S4).

Another exciting avenue to expand this line of research is to include mechanism and underlying machinery as additional context for these patterns. We had hoped to include mechanism as a moderator in our analyses, but we were forced to drop it due to lack of data (see Supplemental Methods - Deviations from predetermined plan). During our review of the literature, we found 77 studies that fit our TGP criteria but only explored molecular (gene expression or epigenetics) rather than phenotypic outcomes in response to stress. While some studies were included in this analysis if they had corresponding trait data, we were unable to incorporate complex molecular datasets into our analysis despite this being an active area of research in the plasticity literature. We believe studies exploring molecular responses and mechanisms are an exciting and informative future direction in the plasticity literature, but recommend studies incorporate some kind of trait-based phenotype along with their molecular data to allow for comparisons across the literature.

Overall, we were interested in exploring how plasticity interactions could inform our understanding of how organisms respond to environmental variability and help broaden our understanding of acclimation and adaptation mechanisms. While we were able to look across a wide variety of systems, stressors, and traits, it is clear that context was critically important and defined what systems we may or may not expect to observe these forms of plasticity. More studies in some of our less represented categories (plants, infection as a stressor, studies following Design B) would be informative moving forward as we may have failed to detect patterns due to small sample sizes. During our systematic review, we also found a diversity in the natural realism of the studies that makes the results more or less applicable to real world predictions. For example, we found a subset of studies that used multiple wild populations or selection lines to compare patterns across evolutionary histories or used stressor amounts or levels based on real values measured in the wild. We encourage the field to incorporate these elements in their study designs whenever feasible to make results as robust and realistic as possible.

## Conclusion

Overall, we found that the effects of transgenerational and developmental plasticity were subtle and highly context dependent, aligning with previous analyses in the plasticity literature. We found evidence for parental effects, but only weak evidence for interactions between parental and offspring developmental plasticity. The timing of parental stress exposure, however, proved important in determining the strength and direction of offspring responses. Plasticity patterns were highly context dependent, both experimentally and biologically, indicating that we should expect stronger plasticity in some species and in certain systems than others. Moving forward, we encourage empiricists to continue exploring these topics in a variety of systems using robust and fully factorial experimental designs and consider incorporating molecular or mechanistic data and/or environmental realism to help better inform our predictions about evolutionary outcomes and species’ persistence in the face of global change.

## Supporting information

Supplemental Results

## Acknowledgements

We would like to thank Wissam Jawad, Suzie Heneghan, and Kirsten Adams for all of their help and efforts in the process of paper screening. Portions of this research were conducted with high performance computing resources provided by Louisiana State University (http://www.hpc.lsu.edu). IPN was supported by an NSF Postdoctoral Research Fellowship in Biology 2305966, ALS was supported by NSF DBE 2216631, and MWK and RV were supported by NSF IOS 2154283 with further support for MWK from Sea Grant NA18OAR4170098.

## Conflict of Interest

The authors declare no conflict of interest for this study.

## Author contributions

IPN and MWK conceived the ideas and designed methodology; IPN, ALS, RV, and CG collected the data; IPN analysed the data; IPN led the writing of the manuscript. All authors contributed critically to the drafts and gave final approval for publication.

## Statement of inclusion

Our study was a systematic review and a meta-analysis of already published data. We strived to collect a representative sample of existing studies and did not limit our search to a particular region or system. Our international authorship team included members with native languages other than English and multiple members fluent in languages other than English, which allowed us to screen a number of studies written in local languages to try and be as inclusive as possible.

## Data availability statement

Data and code are available publicly from the Dryad Digital Repository [DOI: 10.5061/dryad.d2547d8bw]. All studies included in the analysis are listed in the Data Sources section of our references.

## DATA SOURCES

## Design A (Parental and early-life offspring stress and their interaction)

## Design B (Timing of parental stress and transgenerational carryover)

## Notes

https://datadryad.org/dataset/doi:10.5061/dryad.d2547d8bw

### Competing Interest Statement

The authors have declared no competing interest.

