## Supplemental Methods for "Context matters: A meta-analysis of the variable impact of transgenerational and developmental plasticity on responses to stress"

**Supplementary Methods**

***Literature search and study selection***

We accessed Web of Science and Scopus on April 17, 2024. We attempted to screen as much as possible and excluded studies at this stage that involved humans (health, medicine, psychology), agricultural crossbreeding experiments (animal model maternal effects), and purely genetic studies (e.g., maternal effect mutations) although additional screening was required (see first screening details below).

**Web of Science Core Collection:**

*Initial string used:*
**("trans-generation*" OR "transgeneration*" OR "multigeneration*" OR "multi-generation*" OR "intergeneration*" OR "inter-generation*" OR "cross-generation*" OR "non-genetic inheritance" OR "nongenetic inheritance" OR "maternal effect" OR "paternal effect " OR "parental effect") (Topic) not (human* OR child* OR psych* OR patient OR socio* OR econom* OR medic* OR clinical) (Topic)**
=> produced 20,721 studies

Further filtered by:

*Document type (keep):* Article, Proceeding Paper, Early Access, Correction, Letter, Data paper
=> 16,885 studies

*Web of Science Categories (keep)*: Ecology, Environmental Sciences, Genetics Heredity, Evolutionary Biology, Multidisciplinary Sciences, Toxicology, Biochemistry Molecular Biology, Biology, Plant Sciences, Zoology, Marine Freshwater Biology, Entomology, Cell Biology, Developmental Biology, Environmental Studies, Endocrinology Metabolism, Behavioral Sciences, Neurosciences, Reproductive Biology, Fisheries, Physiology, Biodiversity Conservation, Developmental Studies, Oceanography, Microbiology, Forestry, Immunology, Mathematical Computational Biology, Biophysics, Ornithology, Parasitology, Infectious Diseases, Anatomy Morphology, Limnology, Virology, Soil Science, Mycology
=> 6,477 studies

*Publication Titles (exclude):* American Journal of Human Genetics, Diabetes, Human Genetics, Human Heredity, Human Molecular Genetics, Journal of Medical Genetics, Clinical Genetics, European Journal of Human Genetics, Jove Journal of Visualized Experiments, Journal of Environmental Management,
=> 6,348 studies

**Scopus:**

*Initial string used:* ( TITLE-ABS-KEY ( ( "trans-generation*" OR "transgeneration*" OR "multigeneration*" OR "multi-generation*" OR "intergeneration*" OR "inter-generation*" OR "cross-generation*" OR "non-genetic inheritance" OR "nongenetic inheritance" OR "maternal effect" OR "paternal effect " OR "parental effect" ) ) AND NOT TITLE-ABS-KEY ( ( human* OR child* OR psych* OR patient OR socio* OR econom* OR medic* ) ) )
=> Produced 25,452 studies

Further filtered by:

*Document type (keep):* Article, Note, Letter, Data paper, Report
=> 20,488 studies

*Subject Area (keep):* Agricultural and Biological Sciences, Biochemistry Genetics and Molecular Biology, Environmental Science, Pharmacology Toxicology and Pharmaceutics, Multidisciplinary, Immunology and Microbiology, Earth and Planetary Sciences, Neuroscience, Undefined
=> 9,754 studies

*Filter by keyword (exclude):* Cross Breeding, Gene Mutation, Cattle, Genetic Models, Genetic Crosses, Mutation
=> 8,678 studies

***First screening (titles, keywords, abstracts)***

For this first round of screening, we used paper titles, abstracts, and keywords to determine inclusion or rejection. If there was doubt based on these limited data, the study was included in the next round of screening where all records were retrieved and a more detailed reading could be conducted. IPN screened 80% of documents, ALS screened 9%, RV 8%, and the remaining 3% were screened by MWK and two volunteers (KA and WJ).

Rejection Categories:

- Not an empirical study **[Review]**
    - Theoretical, modeling, perspective/opinion piece, review, meta-analysis, methods paper
- Medical/human **[Human]**

    - Study using humans
    - Stressor or traits measure are directly linked to human health concerns without a good parallel in wild systems (i.e., obesity, cardiac health, antibiotic resistance, depression, HIV, alcoholism, substance abuse, diabetes, cancer/tumors, ingredient safety in human products (e.g., palm oil, medications, cocoa powder, hair dye, MSG)) => health/human context
- Agricultural/Veterinary **[Ag/Vet]**
    - Uses a domesticated species used for production (chicken, cattle, pigs, goats; rice, wheat, soybean) or for captive breeding/zoos
    - Stressors or traits that are explicitly in the agriculture/aquaculture/livestock context (i.e., infections that affect fish farming)
    - Uses weird hybrids/strains/cultivars/mutants
    - Test pest traits, but clearly only those related to crop health
- Not applicable trait/stressor/response **[No trait]**
    - Mutation/rates of mutation/epimutation, study uses mutants
    - Measure gene function, epigenetics, microbiome, telomeres
    - Gene manipulation (CRISPR/Cas9), knock downs (RNAi), demethylation (e.g., Azacitidine, inhibition of DNMT)
    - Maternal/paternal/parental effect is inherited gene/genetics (e.g., genes involved in development, gene mutations impacting embryo development or embryogenesis, "maternal-effect genes")
    - A mechanism paper
    - NOTE: Special/separate rejection category for transcriptome/gene expression studies so we can get a count (see below)
    - Sex or sex ratios NOT considered traits
    - *Is there a trait measure of offspring phenotype?* => No => Reject
- Not a TGP study **[Not TGP]**
    - Selection experiment with many generations
    - Breeding study (measuring outcomes of genetic crosses or hybrids)
    - No parental treatments/manipulations
    - Only fecundity, fruit/seed size, egg size or other parental reproductive traits measured but no traits beyond this first stage in offspring, trait measured in offspring is sex or sex ratios
    - Animal model/genetic variance study that calculates “maternal effect” without measuring traits or manipulating parental treatments (be careful, some of these studies may be usable, read the abstract!)
    - Parental size/condition/sex/breeding strategy/age/brood number or time of laying/mate choice are not considered manipulations to parents unless explicitly linked to an environmental context or stressor
    - *Is there manipulation of the parental environment?* => No => Reject

- Other **[Other]**
    - Studying bacteria/microbes/yeast (No prokaryotes, single cell)
    - Not a biological study (culture, economics, geology, humanities)
    - Phylogenetic/hybridization/speciation study
    - Population dynamics, other studies for ecological purposes, like to know the if adults and juveniles live together, temporal changes on the population, description of migration habits (multi-generational migration)
    - Isotope labeling/marking (esp. of otoliths in fish)
    - Fisheries stock assessments

We also wanted to keep track of the studies that included molecular measures and mechanisms as part of the literature search (not a part of the formal meta-analysis).

- Transcriptomics/Gene expression (no phenotypic data) **[Expression]**
    - (Want a rough count of the number of these to report in paper)
    - ****Only keep paper in this category if it satisfies the other keep criteria**
- Methylation/Epigenetics (no phenotypic data) **[Methylation]**
    - (Want a rough count of the number of these to report in paper)
    - ****Only keep paper in this category if it satisfies the other keep criteria**
- Expression/Methylation paper but references another paper with trait data **[Reject - Expression + Trait other paper]**
    - Just want a count, not to be in the Keep category

Keep categories:

- Keep (natural stressors)
- Keep-chemical (lab chemicals, pesticides, pollutants, heavy metals, micro/nano-plastics)
    - Note: These were ultimately not included in our final analyses
- Keep-Trait + Expression (studies we are keeping that have expression/methylation data AND trait data)
    - These were included if they had natural stressors

***Second screening (full records)***

We pulled all records for our second screening and carefully read through methods sections to ensure all papers met our inclusion criteria. All papers had at least two readers who confirmed the choice of inclusion or rejection. Screening at this stage also included dividing studies into either Design A (transgenerational vs. developmental plasticity effect in offspring), Design B (effect of the timing of stress in parents on offspring), or both. For the initial screening, IPN screened 34%, ALS screened 6%, CG screened 28%, RV screened 8%, two undergraduate volunteers (AK and SH) screened 19%, and MWK screened 3%. All papers that moved onto the data mining phase were additionally screened and examined. The rejection category denoting a lack of developmental plasticity ("No Devo") was critical for our study, but also the category that most often required a second opinion during the review process. We define "developmental plasticity" for Design A as a stressor applied to the offspring at one life stage with a trait measured at a later life stage or with some time passing between treatment and measurement as long as all offspring were measured in the same environment.

Rejection Criteria:

- Does not fit one of the designs we want [***See Fig. 1A and 2A and Fig. S2, S3, S4]***
    - Is not fully factorial in either the parent or offspring treatments **[Not factorial]**
    - Does not manipulate or have a clear parental treatment **[No parent treatment]**
    - Does not manipulate offspring treatment **[No offspring treatment]**
    - Does not measure a trait in the offspring (Note: do NOT need measures from parents or early-life) **[Not offspring trait]**
    - Does not use the same stressor in parents and offspring **[Not one stress parent/offspring]**
    - The stress occurs for multiple generations without a factorial design (i.e., a selection experiment) **[Selection]**
    - Does not have a treatment that occurs in an earlier life phase of the offspring with a subsequent measure of traits **[No devo]**

***Data extraction***

Data extractions were performed by IPN (46%), ALS (16%), RV (13%), and CG (16%). IPN, CG, and ALS were responsible for metaDigitising the data. All data were double checked for accuracy by IPN. 19 papers were missing or did not report the necessary data for extraction. ALS contacted the corresponding authors and requested raw data for each of these papers. We received data from eight authors and these studies are included in the analysis. Another two papers were only missing error data and so we were able to impute standard deviations and retain these studies (see Methods). This left nine studies where we were unable to retrieve enough data to include in the analysis despite them passing our eligibility criteria.

### ***Deviations from predetermined analysis plans***

We were able to follow our predetermined plan for analyses as outlined in our Methods and Supplemental Methods with few deviations. One moderator we planned to include but were not able to due to lack of information within studies was the mechanism of transfer (e.g., hormones, epigenetics, maternal provisioning). We do believe this would be a worthwhile endeavor in the future. We also collapsed our stressor categories down to the five (presented in the manuscript) to ensure that we had a large enough sample size in each category. In particular, resource quality encompassed stressors such as diet quality and season/photoperiod while species interactions absorbed predation/herbivory, competition, and social interactions. We also collapsed annual and perennial plants into one category "plant" in our taxonomy moderator due to small sample sizes. For trait type, we followed the categories outlined by Yin et al. (2019) but decided to include "Behavior" as an additional category.
