## Supplemental Results for "Context matters: A meta-analysis of the variable impact of transgenerational and developmental plasticity on responses to stress"

**SUPPLEMENTAL FIGURES**

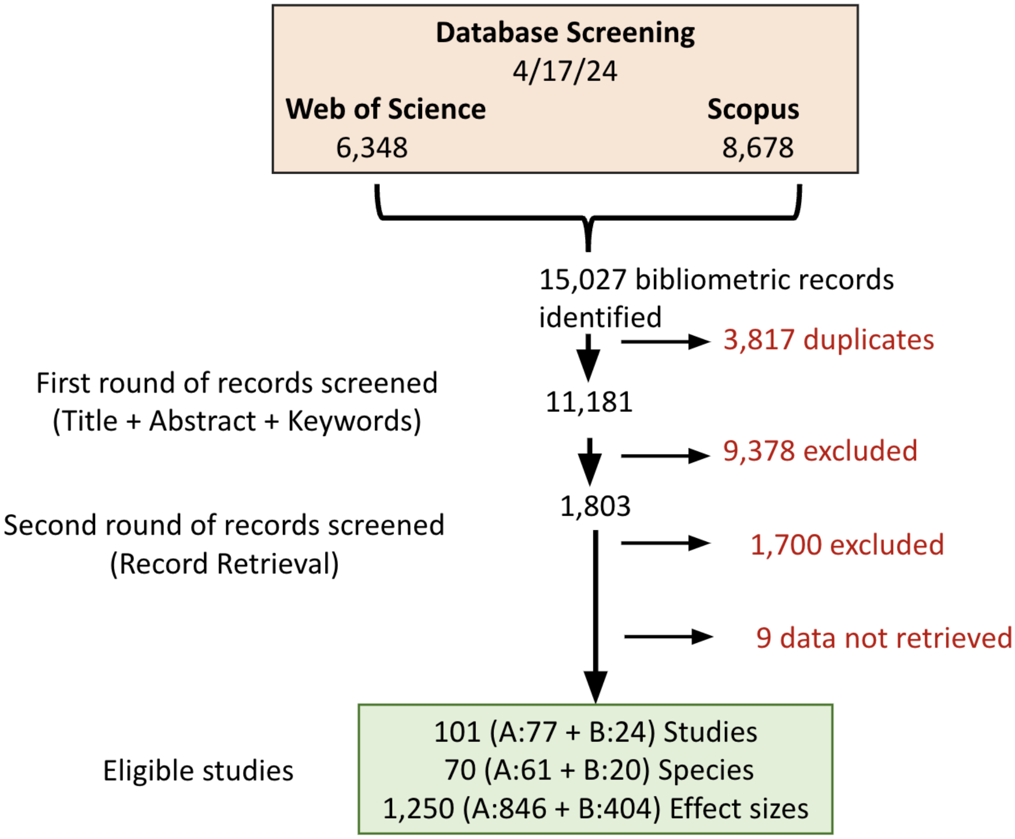


**Fig. S1** Systematic literature review flow chart (PRISMA) illustrating the number of studies rejected at each round of screening and the total number of studies, species, and effect sizes remaining and used in our analyses.

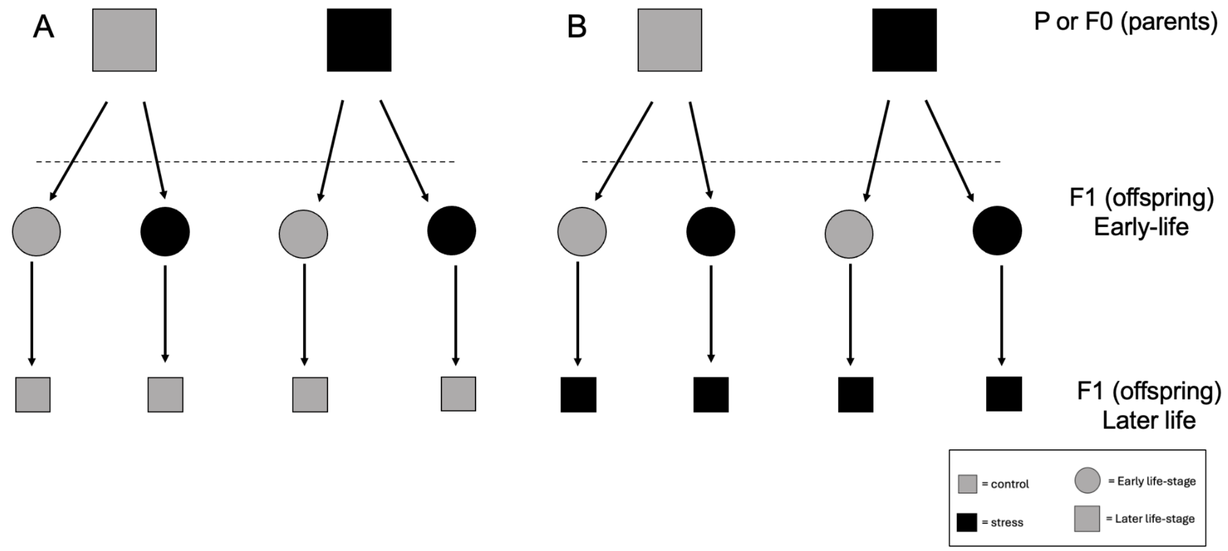


**Fig. S2** Schematics of Design A that do not feature fully factorial designs but were accepted in our meta-analysis.

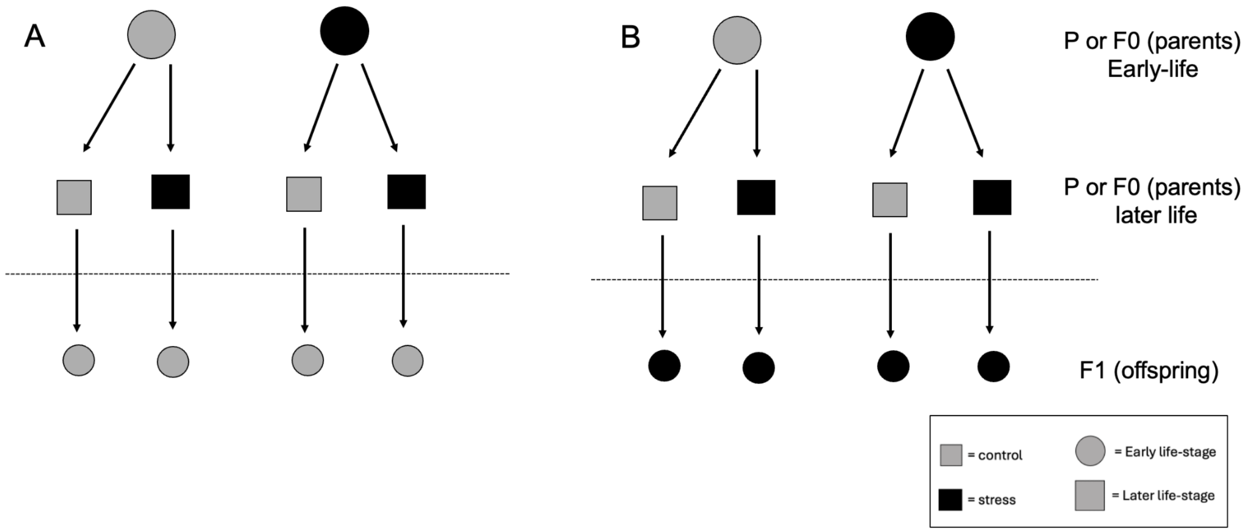


**Fig. S3** Schematics of Design B that do not feature fully factorial designs but were accepted in our meta-analysis.

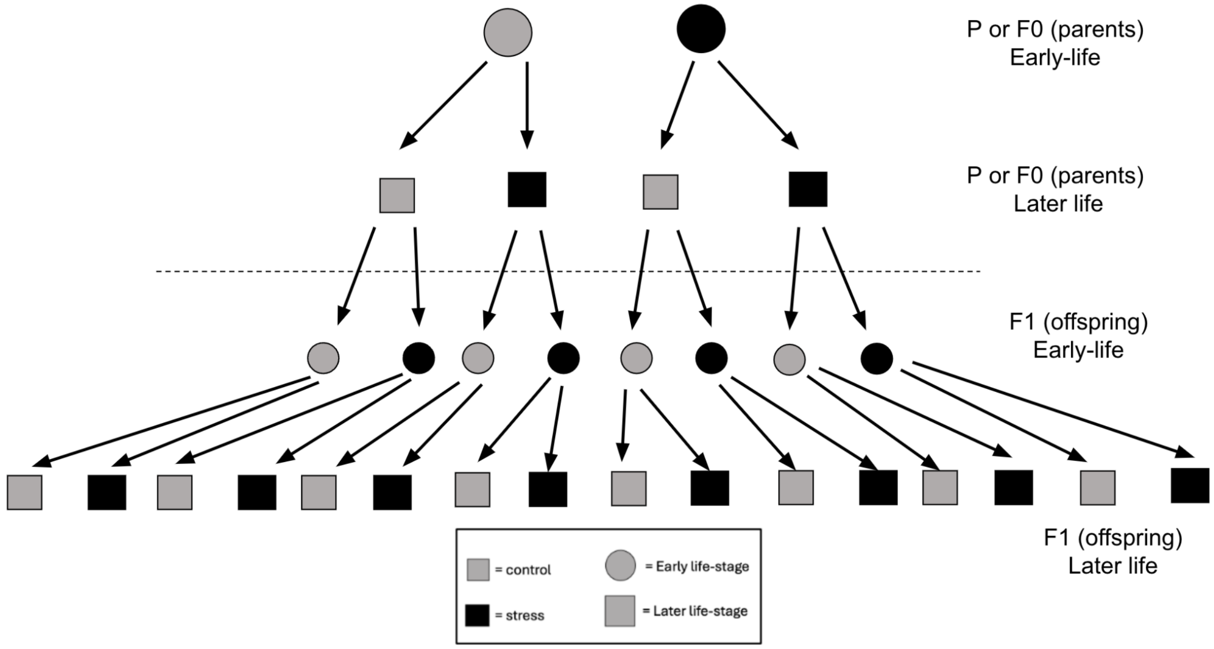


**Fig. S4** Schematic of designs where we were able to collect data for both Design A and B. We argue that this is the most robust design for detecting interactions between various forms of plasticity and should be followed if possible in studies moving forward.

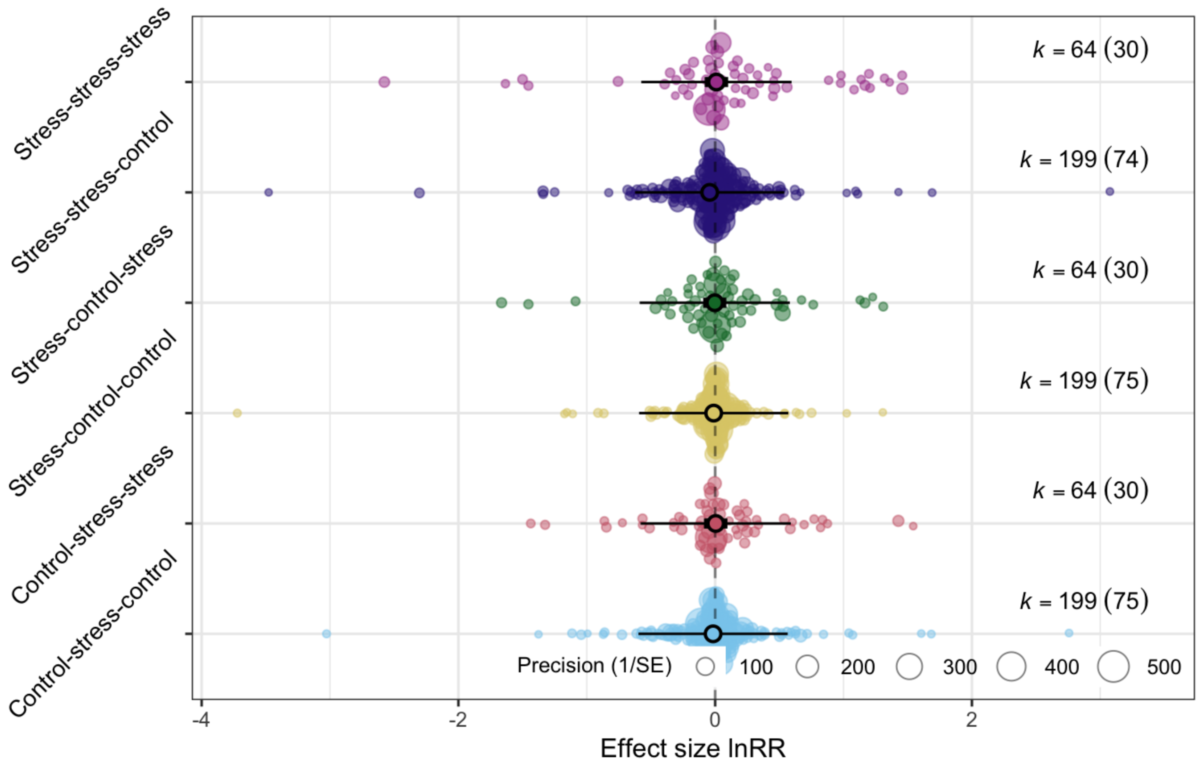


**Fig. S5** Orchard plot showing the effect size estimates for each parent – offspring early – offspring later treatment combinations compared to controls for Design A. Individual colored circles represent effect sizes and are scaled by precision (inverse of standard error).
The central thicker circles indicate the mean estimate with thick error bars representing the 95% confidence intervals and thinner bars representing the prediction intervals. K = number of effect sizes (number of studies).

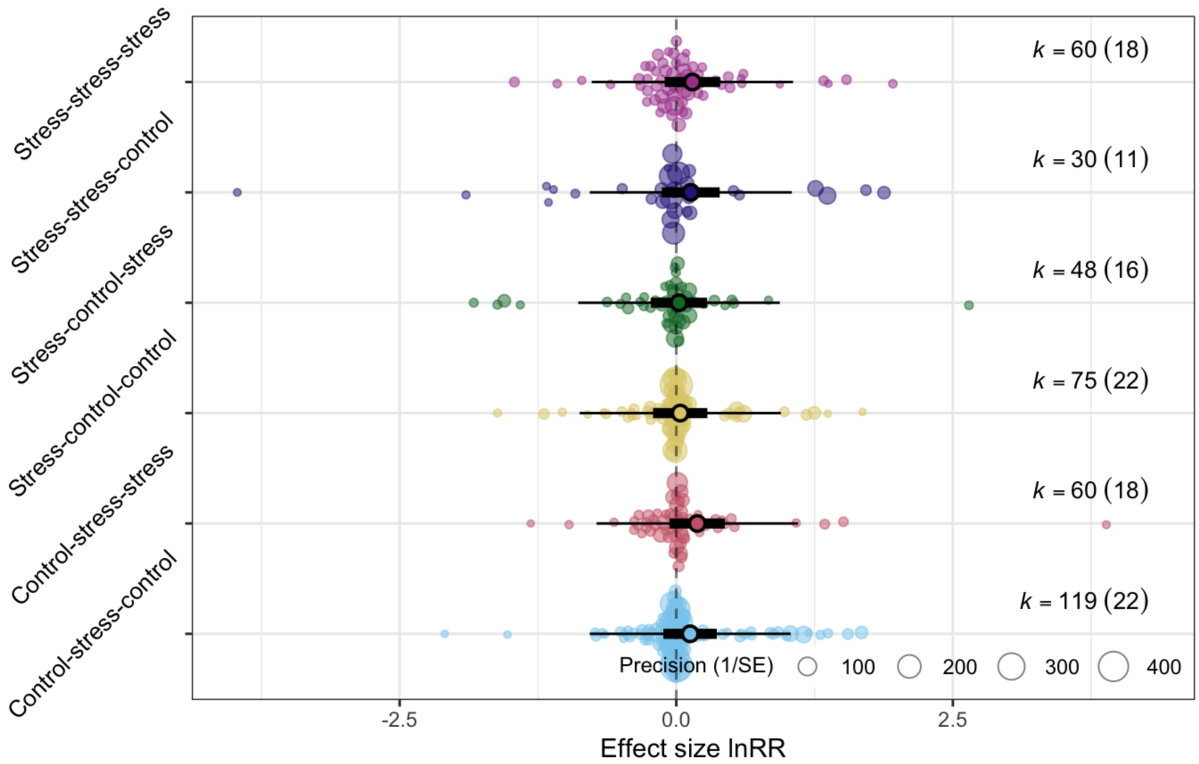


**Fig. S6** Orchard plot showing the effect size estimates for each parent early – parent later – offspring treatment combinations compared to controls for Design A. Individual colored circles represent effect sizes and are scaled by precision (inverse of standard error).
The central thicker circles indicate the mean estimate with thick error bars representing the 95% confidence intervals and thinner bars representing the prediction intervals. K = number of effect sizes (number of studies).

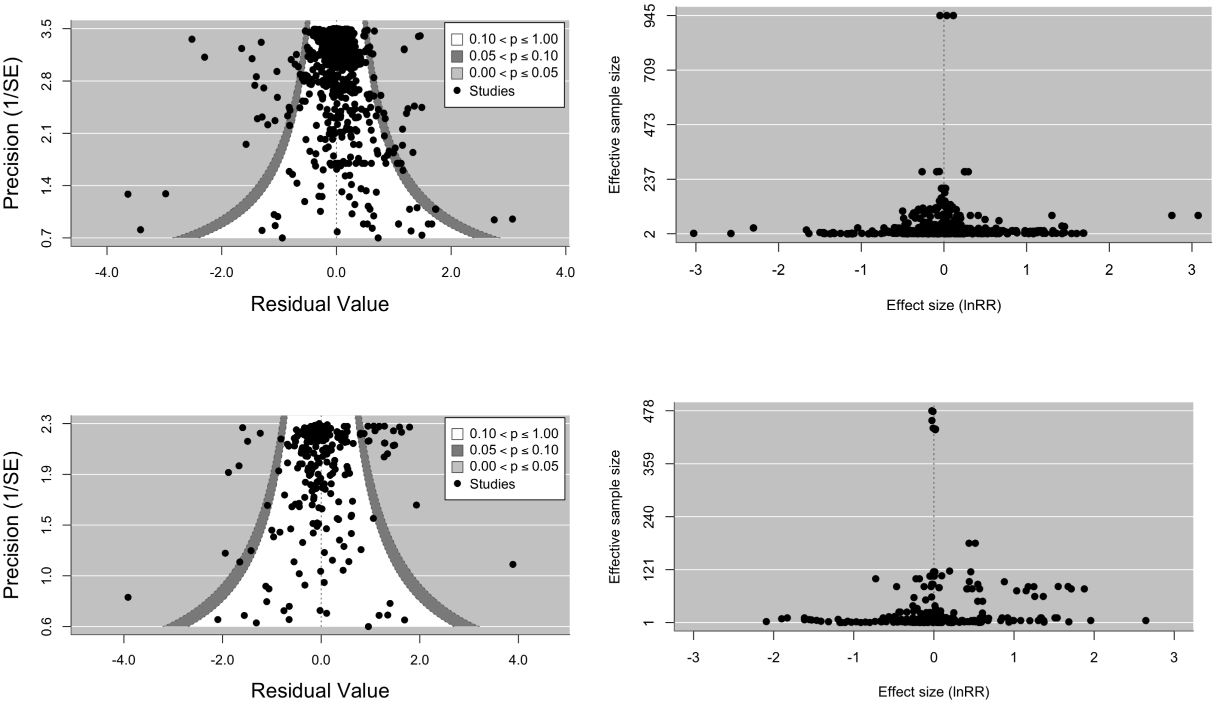


**Fig. S7** Funnel plot graphs for Design A (top row) and Design B (bottom row) to visually inspect evidence of publication bias. The left two graphs show classic funnel plots displaying the distribution of effect sizes (black dots) plotted by their precision (y-axis). The two right graphs show the same data but plotted by effective sample size (y-axis). Symmetrical funnels indicate that there is no evidence of a strong publication bias (that studies with smaller effect sizes are not published).

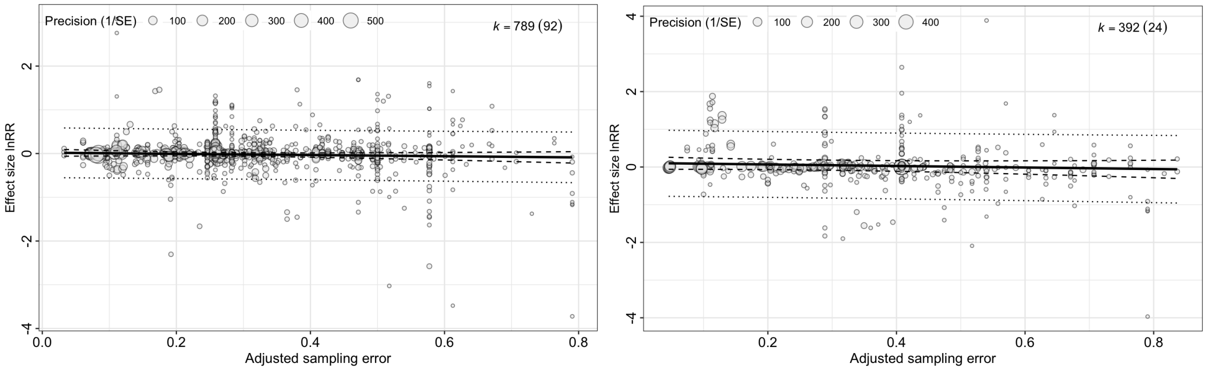


**Fig. S8** Bubble plots to visually determine if there is a small study effect (Design A on the right, Design B on the left). The effect size (calculated as lnRR, y-axis) is plotted against the adjusted sampling error (x-axis) with each circle representing an effect size and the size of the circle representing its precision (inverse of the standard error). K = number of effect sizes (number of studies). The lack of a strong correlation suggests little evidence for small study effects.

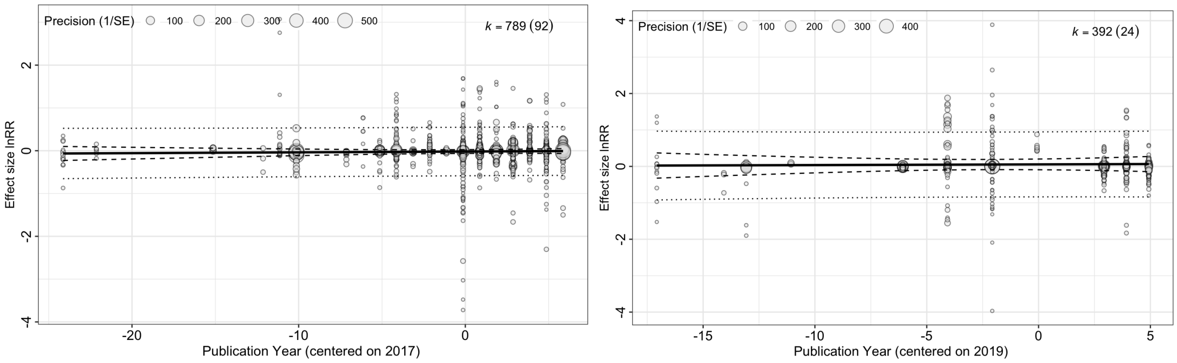


**Fig. S9** Bubble plots to visually examine potential time-lag or decline effects (Design A on the right, Design B on the left). The effect size (calculated as lnRR, y-axis) is plotted against the publication year centered on the mean (x-axis) with each circle representing an effect size and the size of the circle representing its precision (inverse of the standard error). K = number of effect sizes (number of studies). The lack of a strong correlation suggests little evidence for time-lag effects.

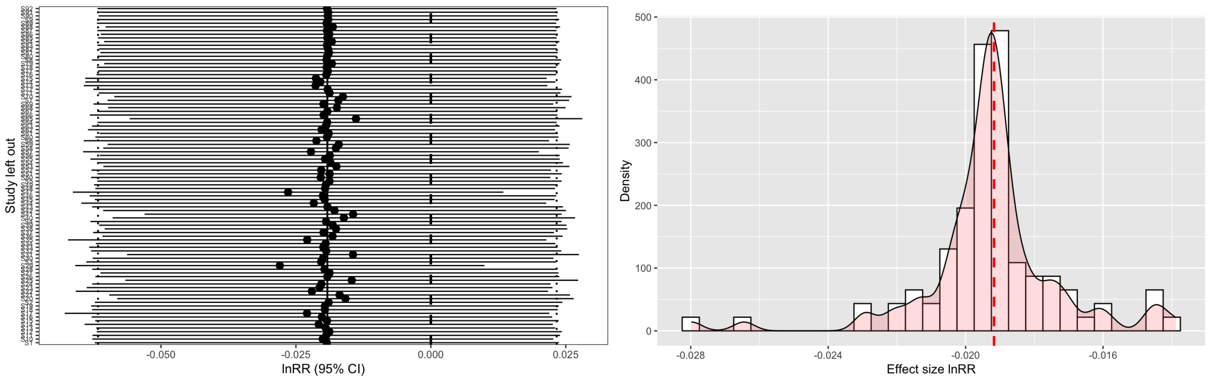


**Fig. S10** Sensitivity analysis graphs for Design A. The left graph shows the results of the leave-one-out analysis where each dot (mean effect size) and error bars (95% confidence intervals) represents the meta-analytic results when an individual study is dropped from the model. The dashed vertical line denotes zero while the dotted lines represent the 95% confidence interval values for the model with all studies included. The right graph shows the same data but as a histogram with the red dashed line representing the mean effect size when all studies are included.

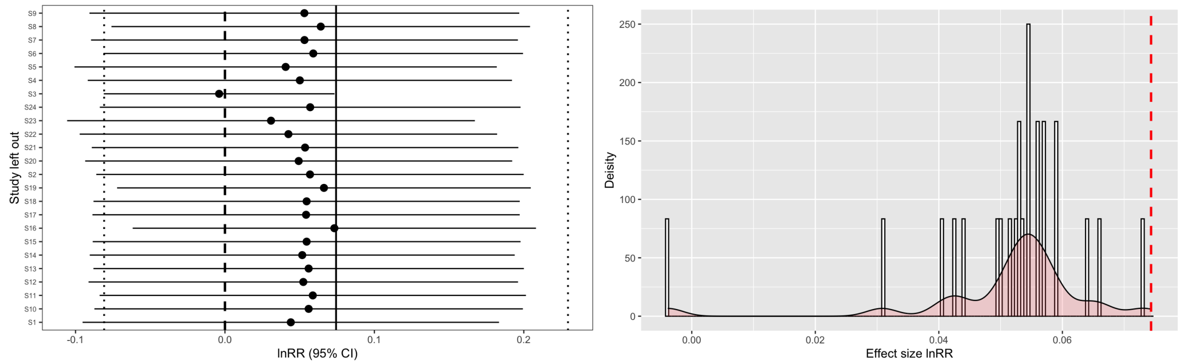


**Fig. S11** Sensitivity analysis graphs for Design B. The left graph shows the results of the leave-one-out analysis where each dot (mean effect size) and error bars (95% confidence intervals) represents the meta-analytic results when an individual study is dropped from the model. The dashed vertical line denotes zero while the dotted lines represent the 95% confidence interval values for the model with all studies included. The right graph shows the same data but as a histogram with the red dashed line representing the mean effect size when all studies are included. Study 3 (S3) appears as a potential outlier and its effects were investigated further.

### SUPPLEMENTAL TABLES

**Table S1** Trait categories within each of the six trait types included in the models for both Design A and Design B with the number of effect sizes per trait category and the number of effect sizes that were associated with a negative fitness correlation and had a sign correction (multiplied by -1) listed in parentheses.

| **Trait type** | **Trait category** | **# of effect sizes (# with sign change) Design A** | **# of effect sizes (# with sign change) Design B** |
|---|---|---|---|
| **Physiological** | | | |
| | Hormone levels | 63 (39) | 15 (15) |
| | Immunity | 27 (6) | 3 (0) |
| | Metabolism/Respiration | 21 (6) | 28 (0) |
| | Speed/movement | 36 (0) | 0 (0) |
| | Thermal tolerance | 87 (0) | 0 (0) |
| | Other | 9 (0) | 0 (0) |
| **Morphological** | | | |
| | Mass/Weight | 87 (6) | 71 (0) |
| | Size/Length | 78 (0) | 80 (0) |
| **Life History** | | | |
| | Rate of maturation/development | 12 (3) | 0 (0) |
| | Development success | 0 (0) | 90 (30) |
| | Size of eggs | 0 (0) | 6 (0) |
| | Survival of offspring | 0 (0) | 12 (0) |
| **Survival** | | | |
| | Survival | 75 (0) | 24 (0) |
| **Reproductive** | | | |
| | Fecundity | 150 (6) | 30 (0) |
| **Behavior** | | | |
| | Behavior | 129 (63) | 0 (0) |
| | Exploratory behavior | 6 (3) | 10 (5) |
| | Feeding | 3 (0) | 11 (0) |
| | Dispersal | 0 (0) | 12 (12) |

**Table S2** Model comparisons between meta-analytic models with different random effect structures in Design A and B to determine best model fit. Phylogeny refers to a correlation matrix that accounts for relatedness of species while Species just refers to the inclusion of the genus and species for each study as a random effect. AIC scores refer to Akaike information criteria, smaller values indicate better model fit.

| **Design A** | **AIC** |
|---|---|
| No random effects | 34261 |
| Study ID | 28559 |
| Study ID, Effect size ID | 612 |
| Study ID, Effect size ID, Species | 595 |
| Study ID, Effect size ID, Phylogeny | 614 |
| **Design B** | **AIC** |
| No random effects | 27307 |
| Study ID | 10772 |
| Study ID, Effect size ID | 482 |
| Study ID, Effect size ID, Species | 481 |
| Study ID, Effect size ID, Phylogeny | 483 |

**Table S3** Results of the meta-analytic intercept model for Design A.

| | **Estimate** | **Standard error** | **t-value** | **Degrees of freedom** | **p-value** | **95% confidence interval (lower)** | **95% confidence interval (upper)** |
|---|---|---|---|---|---|---|---|
| | -0.0192 | 0.0216 | -0.8858 | 788 | 0.376 | -0.0617 | 0.0233 |
| **$I^2$ Total** | 99.824 | **$I^2$ Effect size** | 81.444 | **$I^2$ Study** | >0.0001 | **$I^2$ Species** | 18.379 |

**Table S4** Results from all Design A single moderator meta-regressions fitted with treatment and context moderators. Asterisk next to moderators indicates that results were fitted with models accommodating heteroscedasticity. SE is short for standard error, df for degrees of freedom, CI 95% represents 95% confidence intervals while PI represents predicted intervals. The $R^2$ values shown in parentheses next to each moderator represent marginal $R^2$ values. Bolded p-values indicate significance at $p < 0.05$.

| | Estimate | SE | t-value | df | p-value | CI 95% (lower) | CI 95% (upper) | PI (lower) | PI (upper) |
|---|---|---|---|---|---|---|---|---|---|
| ***Treatment Moderators*** | | | | | | | | | |
| Parent treatment ($R^2$ = 7.1e-5; p = 0.8153) | | | | | | | | | |
| Control | -0.0158 | 0.0261 | -0.6027 | 787 | 0.5469 | -0.0671 | 0.0356 | -0.583 | 0.0356 |
| Stress | -0.0209 | 0.0228 | -0.9139 | 787 | 0.3610 | -0.0657 | 0.0240 | -0.588 | 0.0240 |
| Offspring early-life ($R^2$=0.0001;p =0.7492) | | | | | | | | | |
| Control | -0.0146 | 0.0260 | -0.5607 | 787 | 0.5752 | -0.0656 | 0.0365 | -0.582 | 0.553 |
| Stress | -0.0216 | 0.0229 | -0.9416 | 787 | 0.3467 | -0.0666 | 0.0234 | -0.588 | 0.545 |
| Offspring later life* ($R^2$ =0.003;p= 0.1315) | | | | | | | | | |
| Control | -0.0280 | 0.0154 | -1.8189 | 787 | 0.0693 | -0.0583 | 0.0022 | -0.428 | 0.372 |
| Stress | 0.0174 | 0.0404 | 0.4313 | 787 | 0.6664 | -0.0619 | 0.0967 | -0.874 | 0.909 |
| ***Context Moderators*** | | | | | | | | | |
| Stressor* ($R^2$ = 0.036; **p = 0.0096**) | | | | | | | | | |
| Temperature | 0.0252 | 0.0190 | 1.3289 | 784 | 0.1843 | -0.0120 | 0.0625 | -0.269 | 0.320 |
| Chemistry | 0.0546 | 0.0537 | 1.0160 | 784 | 0.3099 | -0.0509 | 0.1601 | -0.746 | 0.855 |
| **Resource quality** | -0.0679 | 0.0247 | -2.7511 | 784 | **0.0061** | -0.1163 | -0.0194 | -0.521 | 0.385 |
| Species interactions | -0.0843 | 0.0462 | -1.8248 | 784 | 0.0684 | -0.1749 | 0.0064 | -0.863 | 0.694 |
| Infection | -0.0336 | 0.0359 | -0.9367 | 784 | 0.3492 | -0.1041 | 0.0368 | -0.255 | 0.187 |
| Trait* ($R^2$ = 0.026; **p = 0.0024**) | | | | | | | | | |
| Physiology | 0.0250 | 0.0256 | 0.9758 | 783 | 0.3294 | -0.0253 | 0.0754 | -0.292 | 0.342 |
| Morphology | -0.0182 | 0.0253 | -0.7203 | 783 | 0.4715 | -0.0678 | 0.0314 | -0.405 | 0.370 |
| **Life history** | 0.1368 | 0.0484 | 2.8248 | 783 | **0.0049** | 0.0417 | 0.2319 | -0.218 | 0.492 |
| **Survival** | -0.1737 | 0.0741 | -2.3453 | 783 | **0.0193** | -0.3191 | -0.0283 | -0.987 | 0.639 |
| Reproductive | -0.0278 | 0.0432 | -0.6447 | 783 | 0.5193 | -0.1125 | 0.0569 | -0.695 | 0.639 |
| Behavior | -0.0732 | 0.0521 | -1.4055 | 783 | 0.1603 | -0.1756 | 0.0291 | -0.977 | 0.830 |
| Taxonomy ($R^2$ = 0.022; p = 0.2328) | | | | | | | | | |
| Invertebrate | -0.0391 | 0.0285 | -1.3744 | 786 | 0.1697 | -0.0950 | 0.0167 | -0.604 | 0.526 |
| Vertebrate | 0.0306 | 0.0359 | 0.8527 | 786 | 0.3941 | -0.0399 | 0.1011 | -0.536 | 0.597 |
| Plant | -0.0978 | 0.0756 | -1.2928 | 786 | 0.1965 | -0.2463 | 0.0507 | -0.679 | 0.484 |
| Development ($R^2$ = 0.008; p = 0.3385) | | | | | | | | | |
| Biphasic | -0.0399 | 0.0280 | -1.4215 | 787 | 0.1556 | -0.0949 | 0.0152 | -0.610 | 0.530 |
| Direct | 0.0125 | 0.0347 | 0.3587 | 787 | 0.7199 | -0.0557 | 0.0896 | -0.559 | 0.584 |
| Life history traits ($R^2$ = 0.009; p = 0.6946) | | | | | | | | | |
| Internal gestation | 0.0409 | 0.0601 | 0.6803 | 785 | 0.4965 | -0.0772 | 0.1590 | -0.540 | 0.621 |
| External embryos | -0.0327 | 0.0254 | -1.2889 | 785 | 0.1978 | -0.0825 | 0.0171 | -0.603 | 0.538 |
| External gametes | -0.0015 | 0.0648 | -0.0239 | 785 | 0.9809 | -0.1288 | 0.1257 | -0.584 | 0.581 |
| Clonal | 0.0161 | 0.1640 | 0.0979 | 785 | 0.9220 | -0.3058 | 0.3379 | -0.637 | 0.669 |
| Offspring timing ($R^2$ = 0.007; p = 0.531) | | | | | | | | | |
| Hatchling/Germination/Larva | 0.0273 | 0.0618 | 0.4426 | 786 | 0.6582 | -0.0939 | 0.1486 | -0.557 | 0.612 |
| Juvenile/Seedling | -0.0535 | 0.0369 | -1.4510 | 786 | 0.1472 | -0.1258 | 0.0189 | -0.630 | 0.523 |
| Adult | -0.0109 | 0.0280 | -0.3889 | 786 | 0.6975 | -0.0660 | 0.0441 | -0.585 | 0.563 |

**Table S5** Results from Design A meta-regressions fitted with all treatment moderators and their interactions. SE is short for standard error, df for degrees of freedom, CI 95% represents 95% confidence intervals.

| | **Estimate** | **SE** | **t-value** | **df** | **p-value** | **CI95% (lower)** | **CI 95% (upper)** |
|---|---|---|---|---|---|---|---|
| (Intercept) | 0.0102 | 0.0397 | 0.2579 | 783 | 0.7966 | -0.0676 | 0.0881 |
| Parent treatment | -0.0257 | 0.0288 | -0.8931 | 783 | 0.3721 | -0.0821 | 0.0308 |
| Offspring early treatment | -0.0283 | 0.0287 | -0.9832 | 783 | 0.3258 | -0.0847 | 0.0282 |
| Offspring later treatment | -0.0546 | 0.0765 | -0.7139 | 783 | 0.4755 | -0.2047 | 0.0955 |
| Parent * Offspring later | 0.0633 | 0.0609 | 1.0392 | 783 | 0.2990 | -0.0563 | 0.1829 |
| Offspring early * Offspring later | 0.0694 | 0.0606 | 1.1457 | 783 | 0.2523 | -0.0495 | 0.1883 |
| **$R^2$** = 0.005 **Test of Moderators:** F(df1 = 5, df2 = 783) = 0.6246, p-value = 0.6810 | | | | | | | |

**Table S6** Model comparisons with and without accounting for heteroscedasticity for those moderators with high variance of residuals between categories in Design A. AICc scores refer to Akaike information criteria corrected for small sample sizes, smaller values indicate better model fit.

| **Moderator** | | **AICc** |
|---|---|---|
| Offspring later | | |
| | Without | 595 |
| | With | 473 |
| Stressor | | |
| | Without | 591 |
| | With | 445 |
| Trait | | |
| | Without | 595 |
| | With | 332 |

**Table S7** Comparison of Design A meta-regression models incorporating both treatment and context moderators and their interactions. The first (full) model refers to the model that included all moderators and all of their interactions. The second model included all three treatment moderators, but only those context moderators that our model dredging indicated had some explanatory value (stressor category, trait type, taxonomy, developmental trait, life history traits, and timing of trait measures in offspring). Note, that we could not dredge our full model as there were too many moderators, so we ran a series of models and dredged each in multiple rounds to determine which moderators seemed relevant to include in this subsequent model selection approach. We could not accurately determine model structure or weights from these dredging results. The third model included the same suite of treatment and context moderators, but limited interactions to all three treatment moderators and each context moderator (no interactions between context moderators). This third model proved to have the best fit based on AIC, BIC, and AICc scores. Bolded p-values indicate significance at $p < 0.05$.

| | **F statistic** | **p-value** | **$R^2$** | **logLik** | **Deviance** | **AIC** | **BIC** | **AICc** |
|---|---|---|---|---|---|---|---|---|
| 1. Full Model | 1.1292 | 0.1147 | 0.414 | -149.5144 | 299.0289 | 989.0289 | 2404.4116 | 3352.7912 |
| 2. Significant moderators, full interactions | 1.2037 | **0.0379** | 0.472 | -171.2236 | 342.4472 | 849.4472 | 2066.3727 | 1534.2129 |
| 3. Significant moderators, reduced interactions | 1.5432 | **0.0015** | 0.226 | -240.6811 | 481.3622 | 673.3622 | 1109.7158 | 704.4540 |

**Table S8** Results of the meta-analytic intercept model for Design B.

| | **Estimate** | **Standard error** | **t-value** | **Degrees of freedom** | **p-value** | **95% confidence interval (lower)** | **95% confidence interval (upper)** |
|---|---|---|---|---|---|---|---|
| | 0.0744 | 0.0789 | 0.9419 | 391 | 0.3468 | -0.0809 | 0.2296 |
| **$I^2$ Total** | 99.875 | **$I^2$ Effect size** | 46.481 | **$I^2$ Study** | >0.0001 | **$I^2$ Species** | 53.395 |

**Table S9** Results from all Design B single moderator meta-regressions fitted with treatment and context moderators. Asterisk next to moderators indicates that results were fitted with models accommodating heteroscedasticity. SE is short for standard error, df for degrees of freedom, CI 95% represents 95% confidence intervals while PI represents predicted intervals. $R^2$ values represent the $R^2$ marginal values for each moderator. Bolded p-values indicate significance at $p < 0.05$.

| | Estimate | SE | t-value | df | p-value | CI 95% (lower) | CI 95% (upper) | PI (lower) | PI (upper) |
|---|---|---|---|---|---|---|---|---|---|
| ***Treatment Moderators*** | | | | | | | | | |
| Parent early-life ($R^2$ = 0.008; **p = 0.0215**) | | | | | | | | | |
| Control | 0.1215 | 0.0822 | 1.4788 | 390 | 0.1400 | -0.0400 | 0.2831 | -0.793 | 1.036 |
| Stress | 0.0405 | 0.0810 | 0.4995 | 390 | 0.6177 | -0.1188 | 0.1997 | -0.873 | 0.954 |
| Parent later ($R^2$ = 0.014; **p = 0.0015**) | | | | | | | | | |
| Control | -0.0061 | 0.0380 | -0.0731 | 390 | 0.9417 | -0.1692 | 0.1571 | -0.912 | 0.900 |
| Stress | 0.1121 | 0.0799 | 1.4030 | 390 | 0.1614 | -0.0450 | 0.2692 | -0.793 | 1.017 |
| Offspring ($R^2$ = 0.001; p = 0.4048) | | | | | | | | | |
| Control | 0.0814 | 0.7572 | 0.7572 | 390 | 0.4494 | -0.0984 | 0.2216 | -0.858 | 0.981 |
| Stress | 0.0831 | 1.1261 | 1.1261 | 390 | 0.2608 | -0.0698 | 0.2570 | -0.826 | 1.013 |
| ***Context Moderators*** | | | | | | | | | |
| Stressor ($R^2$ = 0.178; **p = 0.0204**) | | | | | | | | | |
| Temperature | -0.2148 | 0.1130 | -1.9010 | 387 | 0.0580 . | -0.4369 | 0.0074 | -1.158 | 0.728 |
| Chemistry | -0.0356 | 0.2620 | -0.1360 | 387 | 0.8919 | -0.5508 | 0.4796 | -1.087 | 1.016 |
| **Resource quality** | 0.2371 | 0.1174 | 2.0201 | 387 | **0.0441** | 0.0063 | 0.4679 | -0.708 | 1.182 |
| Species interactions | 0.2257 | 0.1722 | 1.3105 | 387 | 0.1908 | -0.1129 | 0.5642 | -0.751 | 1.203 |
| Infection | 0.3493 | 0.4807 | 0.7267 | 387 | 0.4678 | -0.5958 | 1.2944 | -0.967 | 1.666 |
| Trait* ($R^2$ = 0.050; p = 0.157) | | | | | | | | | |
| Physiology | -0.0382 | 0.0883 | -0.4325 | 386 | 0.6656 | -0.2118 | 0.1354 | -0.762 | 0.686 |
| Morphology | 0.0808 | 0.0821 | 0.9839 | 386 | 0.3258 | -0.0806 | 0.2422 | -0.750 | 0.922 |
| Life history | 0.1033 | 0.1042 | 0.9910 | 386 | 0.3223 | -0.1016 | 0.3082 | -1.038 | 1.244 |
| Survival | 0.1519 | 0.1011 | 1.5023 | 386 | 0.1338 | -0.0469 | 0.3508 | -0.606 | 0.910 |
| Reproductive | 0.1719 | 0.1011 | 1.7004 | 386 | 0.0899 | -0.0269 | 0.3706 | -0.519 | 0.862 |
| Behavior | 0.3041 | 0.3807 | 0.7988 | 386 | 0.4249 | -0.4444 | 1.0527 | -2.250 | 2.858 |
| Taxonomy* ($R^2$ = 0.12; p = 0.2760) | | | | | | | | | |
| Invertebrate | 0.0922 | 0.1026 | 0.8985 | 18 | 0.4162 | -0.1234 | 0.3078 | -0.778 | 0.963 |
| Vertebrate | -0.0766 | 0.1262 | -0.6068 | 18 | 0.5463 | -0.3416 | 0.1885 | -0.835 | 0.682 |
| Plant | 0.4052 | 0.2463 | 1.6454 | 18 | 0.0474 | -0.1122 | 0.9226 | -1.191 | 2.001 |
| Development* ($R^2$ = 0.040; p = 0.3422) | | | | | | | | | |
| Biphasic | -0.0189 | 0.1125 | -0.1682 | 19 | 0.8682 | -0.2543 | 0.2165 | -0.863 | 0.825 |
| Direct | 0.1551 | 0.1036 | 1.4975 | 19 | 0.1507 | -0.0617 | 0.3720 | -0.930 | 1.240 |
| Life history traits* ($R^2$=0.049; p =0.6197) | | | | | | | | | |
| Internal gestation | -0.0010 | 0.2024 | -0.0051 | 18 | 0.9960 | -0.4263 | 0.4242 | -0.836 | 0.834 |
| External embryos | 0.0434 | 0.0953 | 0.4550 | 18 | 0.6545 | -0.1568 | 0.2435 | -0.837 | 0.924 |
| External gametes | 0.2586 | 0.2037 | 1.2691 | 18 | 0.2206 | -0.1695 | 0.6866 | -1.145 | 1.662 |
| Offspring timing ($R^2$ = 0.147; p = 0.0618) | | | | | | | | | |
| Embryo/Egg/Seed | -0.1437 | 0.1619 | -0.8873 | 388 | 0.3755 | -0.4620 | 0.1747 | -1.053 | 0.766 |
| Hatchling/Germination/Larva | 0.0938 | 0.0961 | 0.9759 | 388 | 0.3297 | -0.0952 | 0.2828 | -0.779 | 0.967 |
| Juvenile/Seedling | -0.0596 | 0.1139 | -0.5233 | 388 | 0.6010 | -0.2834 | 0.1643 | -0.941 | 0.821 |
| **Adult** | 0.3882 | 0.1688 | 2.2995 | 388 | **0.0020** | 0.0563 | 0.7201 | -0.526 | 1.302 |

**Table S10** Results from Design B meta-regressions fitted with all treatment moderators and their interactions. SE is short for standard error, df for degrees of freedom, CI 95% represents 95% confidence intervals. Bolded p-values indicate significance at $p < 0.05$.

| | **Estimate** | **SE** | **t-value** | **df** | **p-value** | **CI95% (lower)** | **CI 95% (upper)** |
|---|---|---|---|---|---|---|---|
| (Intercept) | 0.0491 | 0.1149 | 0.4277 | 386 | 0.6691 | -0.1768 | 0.2751 |
| Offspring treatment | -0.0275 | 0.1136 | -0.2423 | 386 | 0.8087 | -0.2509 | 0.1958 |
| Parent later life treatment | 0.0522 | 0.0763 | 0.6841 | 386 | 0.4944 | -0.0978 | 0.2021 |
| Parent early-life treatment | -0.0432 | 0.0742 | -0.5816 | 386 | 0.5612 | -0.1891 | 0.1027 |
| Parent later * Offspring | 0.0896 | 0.1004 | 0.8924 | 386 | 0.3727 | -0.1078 | 0.2870 |
| Parent early * Offspring | 0.0036 | 0.0968 | 0.0375 | 386 | 0.9701 | -0.1867 | 0.1940 |
| $R^2$ = 0.018 **Test of Moderators:** F(df1 = 5, df2 = 386) = 2.5102, p-value = **0.0297** | | | | | | | |

**Table S11** Model comparisons with and without accounting for heteroscedasticity for those moderators with high variance of residuals between categories in Design B. AICc scores refer to Akaike information criteria corrected for small sample sizes, smaller values indicate better model fit.

| **Moderator** | | **AICc** |
|---|---|---|
| Taxonomy | | |
| | Without | 473 |
| | With | 373 |
| Transmission | | |
| | Without | 478 |
| | With | 455 |
| Development | | |
| | Without | 476 |
| | With | 376 |
| Life history | | |
| | Without | 476 |
| | With | 378 |
| Trait | | |
| | Without | 476 |
| | With | 377 |

**Table S12** Comparison of Design B meta-regression models incorporating both treatment and context moderators and their interactions. The first (full) model refers to the model that included all moderators and all of their interactions. The second model included all three treatment moderators, but only those context moderators that our model dredging indicated had some explanatory value (stressor category, taxonomy, developmental trait, and timing of trait measures in offspring). Note, that we could not dredge our full model as there were too many moderators, so we ran a series of models and dredged each in multiple rounds to determine which moderators seemed relevant to include in this subsequent model selection approach. We could not accurately determine model structure or weights from these dredging results. The third model included the same suite of treatment and context moderators, but limited interactions to the three treatment moderators and each context moderator (no interactions between context moderators). This third model proved to have the best fit based on BIC and AICc scores.

| | **F statistic** | **p-value** | **$R^2$** | **logLik** | **Deviance** | **AIC** | **BIC** | **AICc** |
|---|---|---|---|---|---|---|---|---|
| 1. Full Model | 2.9857 | < 0.001 | 0.709 | -107.9090 | 215.8179 | 549.8179 | 1122.5186 | 1485.0179 |
| 2. Significant moderators, full interactions | 2.4058 | <0.001 | 0.509 | -142.1685 | 284.3369 | 462.3369 | 793.7360 | 536.5036 |
| 3. Significant moderators, reduced interactions | 2.4079 | <0.001 | 0.404 | -168.1824 | 336.3648 | 462.3648 | 702.0883 | 492.4544 |

**Table S13** Results from models incorporating adjusted sampling error to explore potential publication bias for Design A and B. The results remain qualitatively the same as our meta-analytic models (Table S2 and S7) with the intercept remaining non-significant suggesting our datasets are robust to publication bias. SE is short for standard error, df for degrees of freedom, CI 95% represents 95% confidence intervals.

| **Design A** | **Estimate** | **SE** | **t-value** | **df** | **p-value** | **CI 95% (lower)** | **CI 95% (upper)** |
|---|---|---|---|---|---|---|---|
| Intercept | 0.0093 | 0.0275 | 0.3372 | 787 | 0.7361 | -0.0448 | 0.0633 |
| Adjusted sampling error | -0.2752 | 0.1693 | -1.6258 | 787 | 0.1044 | -0.6076 | 0.0571 |

| **Design B** | **Estimate** | **SE** | **t-value** | **df** | **p-value** | **CI 95% (lower)** | **CI 95% (upper)** |
|---|---|---|---|---|---|---|---|
| Intercept | 0.0721 | 0.0719 | 1.0018 | 390 | 0.3171 | -0.0694 | 0.2135 |
| Adjusted sampling error | -0.2029 | 0.2064 | -0.9834 | 390 | 0.3260 | -0.6087 | 0.2028 |

**Table S14** Results from models incorporating the uncertainty of effect size as a moderator (the sampling variance based on effective size) to explore potential small study effect bias for Design A and B. The non-significant p-values and confidence intervals over-lapping zero indicate that there is no significant correlation between the effect size and its error. In other words, our datasets are robust to small study effects. SE is short for standard error, df for degrees of freedom, CI 95% represents 95% confidence intervals.

| **Design A** | **Estimate** | **SE** | **t-value** | **df** | **p-value** | **CI 95% (lower)** | **CI 95% (upper)** |
|---|---|---|---|---|---|---|---|
| Intercept | 0.0201 | 0.0418 | 0.4821 | 787 | 0.6299 | -0.0619 | 0.1021 |
| $\sqrt{Sampling\ variance}$ | -0.1357 | 0.1239 | -1.0954 | 787 | 0.2737 | -0.3789 | 0.1075 |

| **Design B** | **Estimate** | **SE** | **t-value** | **df** | **p-value** | **CI 95% (lower)** | **CI 95% (upper)** |
|---|---|---|---|---|---|---|---|
| Intercept | 0.1078 | 0.0857 | 1.2572 | 390 | 0.2094 | -0.0608 | 0.2764 |
| $\sqrt{Sampling\ variance}$ | -0.2011 | 0.1845 | -1.0900 | 390 | 0.2764 | -0.5639 | 0.1616 |

**Table S15** Results from models incorporating the year of publication (centered on the mean) as a moderator to explore the potential for a time-lag or decline bias for Design A and B. The non-significant p-values and confidence intervals over-lapping zero of the regression slope (estimate of year) indicate that there is no significant correlation between the effect size and year and therefore no evidence for time-lag effects. SE is short for standard error, df for degrees of freedom, CI 95% represents 95% confidence intervals.

| **Design A** | **Estimate** | **SE** | **t-value** | **df** | **p-value** | **CI 95% (lower)** | **CI 95% (upper)** |
|---|---|---|---|---|---|---|---|
| Intercept | -0.0189 | 0.0217 | -0.8710 | 787 | 0.3840 | -0.0614 | 0.0237 |
| Year (centered on 2019) | 0.0019 | 0.0033 | 0.5652 | 787 | 0.5721 | -0.0047 | 0.0085 |

| **Design B** | **Estimate** | **SE** | **t-value** | **df** | **p-value** | **CI 95% (lower)** | **CI 95% (upper)** |
|---|---|---|---|---|---|---|---|
| Intercept | 0.0552 | 0.0749 | 0.7371 | 390 | 0.4615 | -0.0920 | 0.2024 |
| Year (centered on 2017) | 0.0019 | 0.0108 | 0.1737 | 390 | 0.8622 | -0.0193 | 0.0230 |

**Table S16** Results of the meta-analytic model using data from Design B, leaving out the potential outlier study (S3).

| | Estimate | Standard error | t-value | Degrees of freedom | p-value | 95% confidence interval (lower) | 95% confidence interval (upper) |
|---|---|---|---|---|---|---|---|
| | -0.0056 | 0.0401 | -0.1405 | 379 | 0.8883 | -0.0844 | 0.0731 |
| **$I^2$ Total** | 99.77 | **$I^2$ Effect size** | 84.088 | **$I^2$ Study** | >0.0001 | **$I^2$ Species** | 15.682 |

**Table S17** Results of the meta-analytic model using data from Design A, leaving out effect sizes that required imputed standard deviation values.

| | Estimate | Standard error | t-value | Degrees of freedom | p-value | 95% confidence interval (lower) | 95% confidence interval (upper) |
|---|---|---|---|---|---|---|---|
| | -0.0202 | 0.0216 | -0.9377 | 755 | 0.3507 | -0.0626 | 0.0222 |
| **$I^2$ Total** | 99.828 | **$I^2$ Effect size** | 81.121 | **$I^2$ Study** | 0.0217 | **$I^2$ Species** | 18.489 |

**Table S18** Results of the meta-analytic model using data from Design A, leaving out effect sizes that required a continuous correction.

| | Estimate | Standard error | t-value | Degrees of freedom | p-value | 95% confidence interval (lower) | 95% confidence interval (upper) |
|---|---|---|---|---|---|---|---|
| | -0.0139 | 0.0200 | -0.6922 | 779 | 0.4890 | -0.0532 | 0.0254 |
| **$I^2$ Total** | 99.795 | **$I^2$ Effect size** | 81.823 | **$I^2$ Study** | 1.160 | **$I^2$ Species** | 16.812 |

**Table S19** Results of the meta-analytic model using data from Design B, leaving out effect sizes that required a continuous correction.

| | Estimate | Standard error | t-value | Degrees of freedom | p-value | 95% confidence interval (lower) | 95% confidence interval (upper) |
|---|---|---|---|---|---|---|---|
| | 0.0688 | 0.0779 | 0.8826 | 370 | 0.3780 | -0.0845 | 0.2220 |

| | | | | | | | |
|---|---|---|---|---|---|---|---|
| **$I^2$ Total** | 99.795 | **$I^2$ Effect size** | 46.847 | **$I^2$ Study** | 19.806 | **$I^2$ Species** | 33.142 |

**Table S20** Results of models using differing values of rho to construct the variance-covariance matrix for Design A. Overall, our results are robust to changes in the value of rho, so we continued with the standard value of 0.5. SE is short for standard error and CI 95% represents 95% confidence intervals.

| Correlation (ρ) | Overall effect | SE | p-value | 95% CI (lower) | 95% CI (upper) |
|---|---|---|---|---|---|
| 0.3 | -0.0213 | 0.0226 | 0.3475 | -0.0657 | 0.0232 |
| 0.5 | -0.0192 | 0.0216 | 0.3760 | -0.0617 | 0.0233 |
| 0.7 | -0.0171 | 0.0214 | 0.4251 | -0.0592 | 0.0250 |
| 0.9 | -0.0144 | 0.0217 | 0.5067 | -0.0571 | 0.0282 |

**Table S21** Results of models using differing values of rho to construct the variance-covariance matrix for Design B. Overall, our results are robust to changes in the value of rho, so we continued with the standard value of 0.5. SE is short for standard error and CI 95% represents 95% confidence intervals.

| Correlation (ρ) | Overall effect | SE | p-value | 95% CI (lower) | 95% CI (upper) |
|---|---|---|---|---|---|
| 0.3 | 0.0485 | 0.0693 | 0.4843 | -0.0877 | 0.1847 |
| 0.5 | 0.0512 | 0.0695 | 0.4619 | -0.0855 | 0.1878 |
| 0.7 | 0.0545 | 0.0701 | 0.4373 | -0.0833 | 0.1924 |
| 0.9 | 0.0581 | 0.0712 | 0.4154 | -0.0820 | 0.1981 |
